# The development of ToF-SIMS for in-situ glycosaminoglycan analysis

**DOI:** 10.64898/2026.05.06.723150

**Authors:** Lorna K. Milne, Jamie L. Thompson, Raina Ramnath, Simon Satchell, Rebecca L. Miller, Lena Kjellén, Kenton P. Arkill, Catherine L. R. Merry, Andrew L. Hook

## Abstract

Glycosaminoglycans (GAGs) are linear polysaccharides with essential roles in a myriad of biological processes. Despite their biological importance, methods to determine both spatial and compositional information is limited. Time-of-flight secondary ion mass spectrometry (ToF-SIMS) provides spatially resolved compositional information of biological molecules without enzymatic digestion or label incorporation, enabling unbiased analysis independent of enzyme or label selectivity, overcoming many current limitations in GAG analysis. Here, we present the identification and validation of GAG discriminatory ions from biological samples by comparison of spectra from purified GAGs and cells with genetically modified GAG biosynthetic pathways. Ions discriminatory of specific GAG sub-families are identified and related to GAG structural components. The analysis is applied to human induced pluripotent stem cells engineered to lack heparan sulphate (HS), where compensatory changes in GAG display that link to function were observed. Furthermore, the broad applicability and spatial resolution of the technique is highlighted through detection of a disease-induced reduction in HS within the individual glomeruli of diabetic mice.

## Main

Glycosaminoglycans (GAGs) are a heterogeneous family of linear polysaccharides consisting of combinations of disaccharide units (Figure 1a). With the exception of hyaluronan (HA), they are synthesised covalently bound to a core protein and are typically displayed at the cell surface or incorporated into the extracellular matrix (ECM) where they mediate interaction with the cell’s local microenvironment^1^. The presence of sulphate motifs largely determines GAG function, where they regulate numerous biological processes including embryonic patterning and vascular permeability^2,3^. Disruption of GAG biosynthesis and composition is a hallmark of several diseases, including cancer, sepsis and diabetes^4–6^.

**Figure 1.**
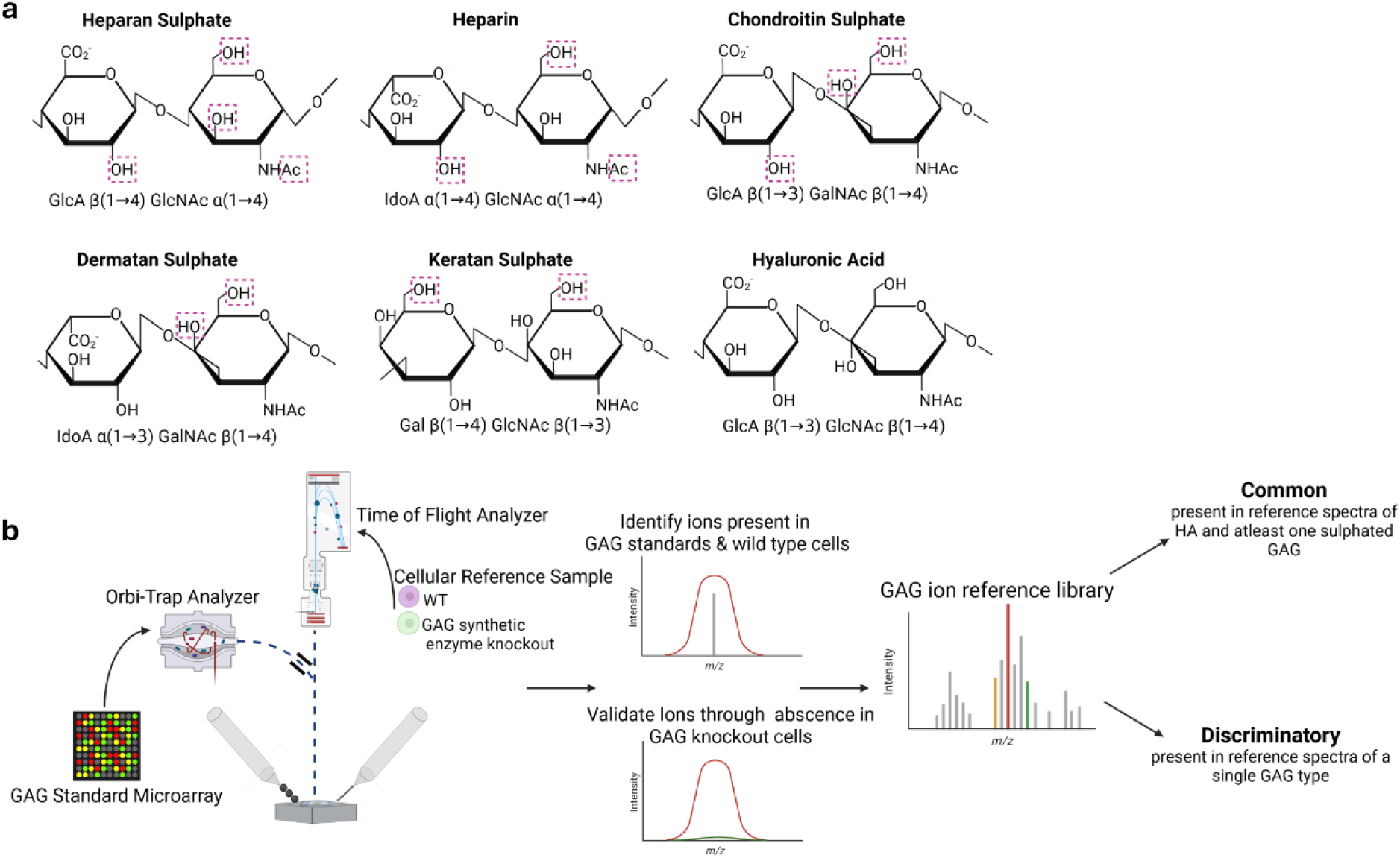
Schematic of GAG structures and workflow used in generation of a GAG ion reference library. **(a)** Disaccharide structures of individual GAG species. GlcA; D-glucuronic acid, IdoA; L-iduronic acid; Gal; D-galactose, GlcNAc; *N*-acetyl- D-glucosamine, GalNAc *N*-acetyl-D-galactosamine. Pink squares indicative of potential sites of sulphation modification **(b)** Orbi-SIMS reference spectra were acquired from a GAG standard microarray as previously described^7^. Ions found in individual GAG reference spectra were identified in wild type cells and validated as GAG discriminatory through absence in GAG biosynthetic knockout cells. Two categories of ions were determined for GAG analysis: common GAG and individual GAG discriminatory ions.

Despite their physiological importance, GAG analysis is complicated by their structural variability, non-template driven synthesis and presence of multiple GAG types and chains on a single core protein. This limits correlation with transcriptomics and underpins an overall lack of understanding of the relationship between GAG composition, position and function. Current methods of GAG analysis typically prioritise either spatial or compositional information. Commercially available antibodies are routinely used for spatial analysis of GAGs; however, GAG-specific probes are limited in comparison to other biomolecules (including other glycans). Further, there is a lack of commercially available antibodies to discriminate between *O-*sulphation motifs within the GAG chain. Lectin-based probes can be used to tag GAGs, however, their ability to recognise a GAG is often non-specific or tissue-type dependent^8^. Disaccharide analysis remains the gold-standard to study GAG composition in a quantitative manner^9,10^. However, the technique is typically dependent on relatively large amounts of starting material and the requirement to digest and remove GAGs from their spatial context. On-sample digestion has been used to complete disaccharide analysis on GAGs extracted from histologically distinct regions in frozen tissue, providing some spatial infomation^11^.

Mass spectrometry imaging (MSI) provides a potential intermediate, combining compositional data through the generation of mass spectra and spatial data through site specific ion generation. Desorption electrospray ionisation (DESI) has been used for the analysis of heparan sulphate (HS) and heparin (HP) octasaccharides containing up to seven sulphate motifs but the technique has not yet been utilised for spatial analysis^12^. The spatial distribution of chondroitin sulphate (CS) has been assessed in the brain using matrix desorption electrospray ionisation (MALDI) to a pixel size of 150 μm^13^. However, its application to GAG analysis remains limited due to the labile nature of sulphates during the ionisation process. A third method of MSI, secondary ion mass spectrometry (SIMS) uses a focused primary ion beam to fragment the upper 1-2 nm of a sample’s surface. The mass of the ions produced is measured using either a time of flight (ToF) or orbi-trap (Orbi) mass analyser^14^. It has previously been utilised for the *in-situ* analysis of lipids^15^, metabolites^14^, proteins^16^ and RNA^17^. SIMS has, in principle, a spatial resolution of down to ≈100 nm^18^. Whilst this is often lower in biological samples, it can still be used to enable single cell analysis^19^. Furthermore, SIMS benefits from being both label and matrix free. It is also untargeted, simultaneously collecting data on all biological molecules at once, enabling a potential high throughput approach to multi-omics with sample rates as high as 1000 pixels/s.

We have previously shown that SIMS can discriminate between nanogram quantities of individual GAG standards, identify HP contamination with over sulphated CS to a specificity of 0.001% (w/w) and discriminate HP from different animal sources^7,20^. However, there is limited use of SIMS for *in-situ* characterisation of GAGs within biological samples. A handful of studies have identified ions likely associated with GAGs generally, but there are no previous reports of SIMS ions that discriminate between GAG type or that can inform on GAG composition^21,22^

In this study we have identified GAG-derived ions within biological samples that can be linked to specific GAG types and potential sulphation motifs (Figure 1b). This was achieved through the identification of GAG specific ions in the SIMS spectra of reference samples, their detection in unmodified cell lines and absence in multiple genetically modified cell lines with altered GAG biosynthesis. Identified ions were then assessed in individual glomeruli of healthy and diabetic mice to test whether ions could be used to provide concomitant spatial and compositional information about GAGs from complex pathological tissue.

## Results

### Identification of potential GAG associated ions from biological samples

ToF-SIMS spectra were acquired for wild type (WT) human induced pluripotent stem cells (hiPSCs) that had not undergone any method of fixation or freezing prior to analysis and compared with Orbi-SIMS reference spectra of purified GAGs^7^. Common GAG ions were identified, defined as those found in the reference spectra of HA and at least one sulphated GAG (supplementary table 2-3) in addition to the ToF-SIMS spectrum acquired from WT hiPSCs. A previously described chemical filter was used to provide possible assignments based upon the chemical structure of GAGs^7^. For HA, sulphate was excluded during assignment. Peaks which could not be assigned to chemical structures are listed as unknown. Putative discriminatory ions could be identified for all GAG types, defined through (i) presence exclusively in the reference spectra of a single GAG type and (ii) detection in WT hiPSCs (supplementary tables 4-8).

### Assessment of sample preparation on GAG ion yield

We compared common methods of SIMS sample preparation to assess their impact on GAG-associated ion yield and retention of cell morphology, both of which are critical for integration into multi-method analytical workflows. Total common GAG ion intensity was compared between all methods of sample preparation. Under all conditions, common GAG ions could be identified in multiple replicates (supplementary figure 1a). Both untreated and chemical fixation methods best retained typical cell morphology when compared to brightfield imaging of cells in culture (supplementary figure 1b/c). Based on typical morphology retention, the detection of putative GAG ions and advantages for sample storage and transportation, chemical fixation was used to prepare all subsequent samples.

### Generation of a HS deficient hiPSC line

Previous reports of SIMS of GAGs in biological samples have centred around the use of sulphate containing species such as SO_4_^-^ which are not unique to GAGs^21,22^. We have previously used SIMS to discriminate between GAG types and animal source species^7,20^. Here, we aimed to identify SIMS ions which define GAG species in complex biological systems. GAG discriminatory ions were defined *in-situ* by comparing SIMS spectra from purified GAGs with those of cellular samples where individual GAGs or specific sulphated motifs were differentially present against an unmodified cellular background, confirmed by well-established methods (supplementary Figures 2 and 3).

To achieve this, we analysed cell lines with knockout/knock-in of enzymes at specific points of the GAG biosynthetic pathway^23^, the spectra of which, when compared with that of WT cells could be used to identify through absence, ions associated with a specific GAG synthesis step. This strategy has previously been used to great success for characterisation of glycan-binding ligands^24^ but has not been applied to identify ions of interest within a SIMS spectrum.

We took advantage of a fully defined, GAG-free culture system available for hiPSC culture and targeted the HS polymerase EXT2 (an essential component of the HS biosynthetic pathway^25^) using CRISPR-Cas9 gene editing to generate a HS-deficient cell line (supplementary methods and supplementary figure 2). In agreement with recent reports, this line could be maintained without additional GAG supplementation, retaining typical hiPSC morphology and marker expression^26^. Absence of HS was confirmed using flow cytometry (supplementary 3ai/aii) and disaccharide analysis (supplementary 3b). To characterise the full sulphated GAG profile of the hiPSCs, the cells were metabolically radiolabelled with [^35^S] sodium sulphate, followed by isolation of ^35^S-labeled GAGs. These were then analysed before and after digestion with GAG-specific enzymes (supplementary figure 3c). The *Ext2^-/-^* cells lacked HS but had relative increases in both CS/DS and a population likely to be predominantly KS due to its resistance to both specific heparinase and chondroitinase ABC digestion, in agreement with relatively strong staining of the *Ext2^-/-^* cells with KS-specific antibodies (supplementary figure 3d).

### ToF-SIMS enables GAG detection in diverse cell types

Common GAG ion intensity was compared between WT hiPSC and those lacking HS synthesis. We also analysed an *Ext1^-/-^* Chinese Hamster Ovary (CHO) line which have previously been described to lack HS synthesis^23^. In all lines, the distribution of total common GAG ion intensity was largely homogenous across the cell surface (Figure 2). Total common GAG ion intensity was significantly reduced in HS knockout hiPSCs (*p=0.047,* Figure 2aii). No significant difference in total common GAG ion intensity was observed between WT and HS knockout CHO cells (Figure 2bii), potentially due to the comparatively low level of GAG synthesis in these cells.

**Figure 2.**
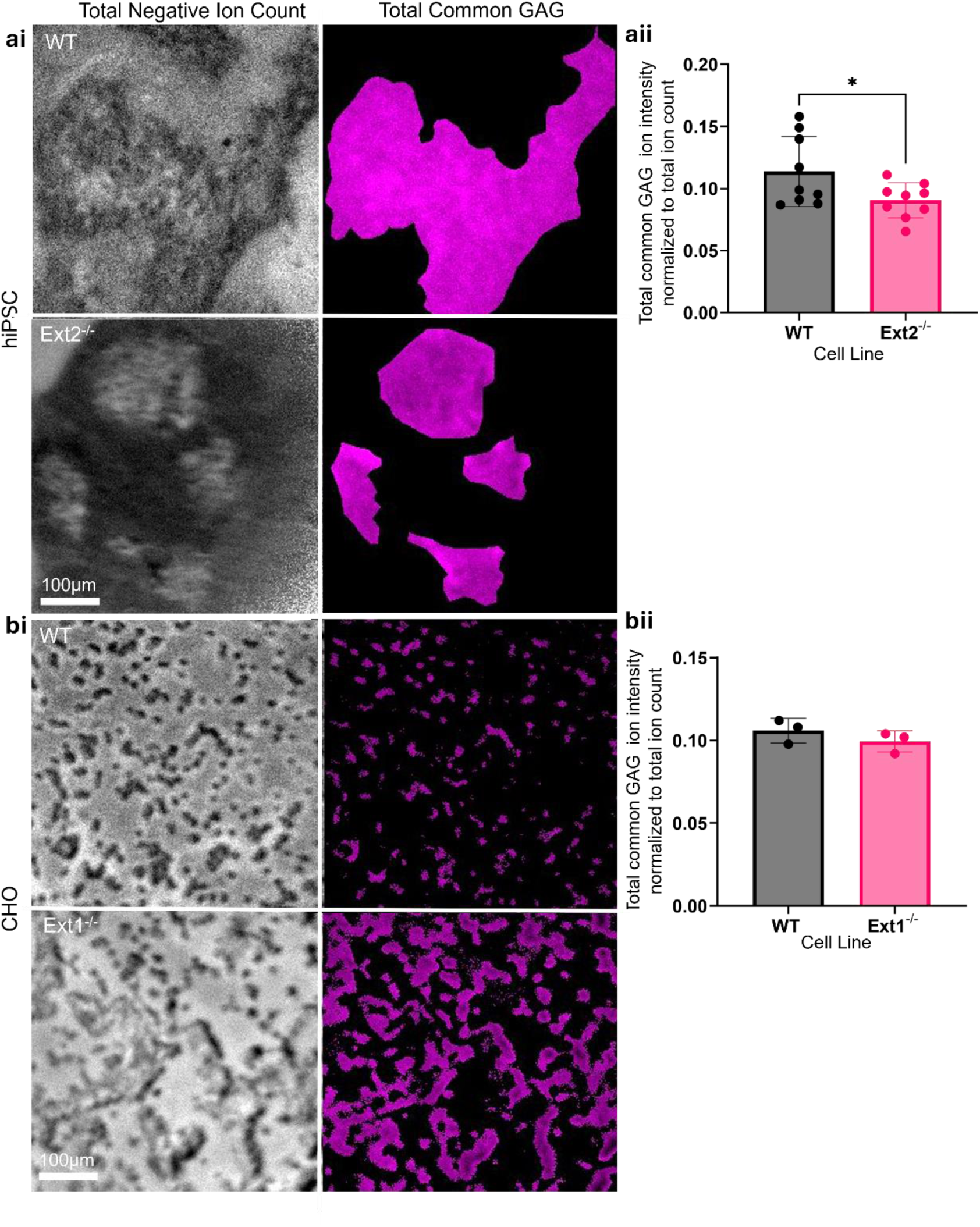
Common GAG ions are detected in multiple cell lines. (ai, bi) Example ToF-SIMS ion images depicting (left) total negative ion count and (right) summed total common GAG ion intensity within a region of interest in (top) WT and (bottom) HS deficient cell lines (ai) *Ext2^-/-^* hiPSCS (bi) *EXT1^-/-^* CHO. Scale bar = 100 μm (aii, bii,) Comparison of total common GAG ion intensity between WT and HS deficient cell lines (aii) *Ext2^-/-^* hiPSCs (n=3 of N=3) (bii) *Ext1^-/-^* CHO (n=3). Error bars show ± SEM. A significant reduction in common GAG ion intensity was observed for *EXT2^-/-^* hiPSCs (p=0.047, unpaired t-test). No significant difference in total common GAG ion intensity was observed between CHO WT and *EXT1^-/-^* lines.

Here we show the development of analytical pipelines for the integration of ToF and Orbi-SIMS spectra and the first report of genetically modified cell lines for the validation of biomolecule discriminatory ions in SIMS, with previous methods utilizing heavy metals or isotopic labelling^27,28^.

### Validation of a defined set of ions to identify HS in complex biological samples

Identification of HS discriminatory ions was based on the criteria: (i) detection exclusively in HS reference spectra (and not in HP), (ii) presence in all WT cell lines and (iii) absence in all HS knockout lines (Figure 3bi/ii, ci/ii, supplementary figure 4). Following the validation process, nine ions were identified that fulfilled these criteria and were therefore categorised as HS discriminatory ions. The absence of these ions was statistically significant across all knockout lines *(p=<0.0001* in all HS knockout lines).

To further confirm these ions as HS discriminatory, their intensity was assessed in a *Chsy1^-/-^* CHO line that is CS/DS deficient, but where HS synthesis is retained. No significant difference in total HS ion intensity was observed, confirming that these ions are HS discriminatory *in-situ* (Figure 3d).

**Figure 3.**
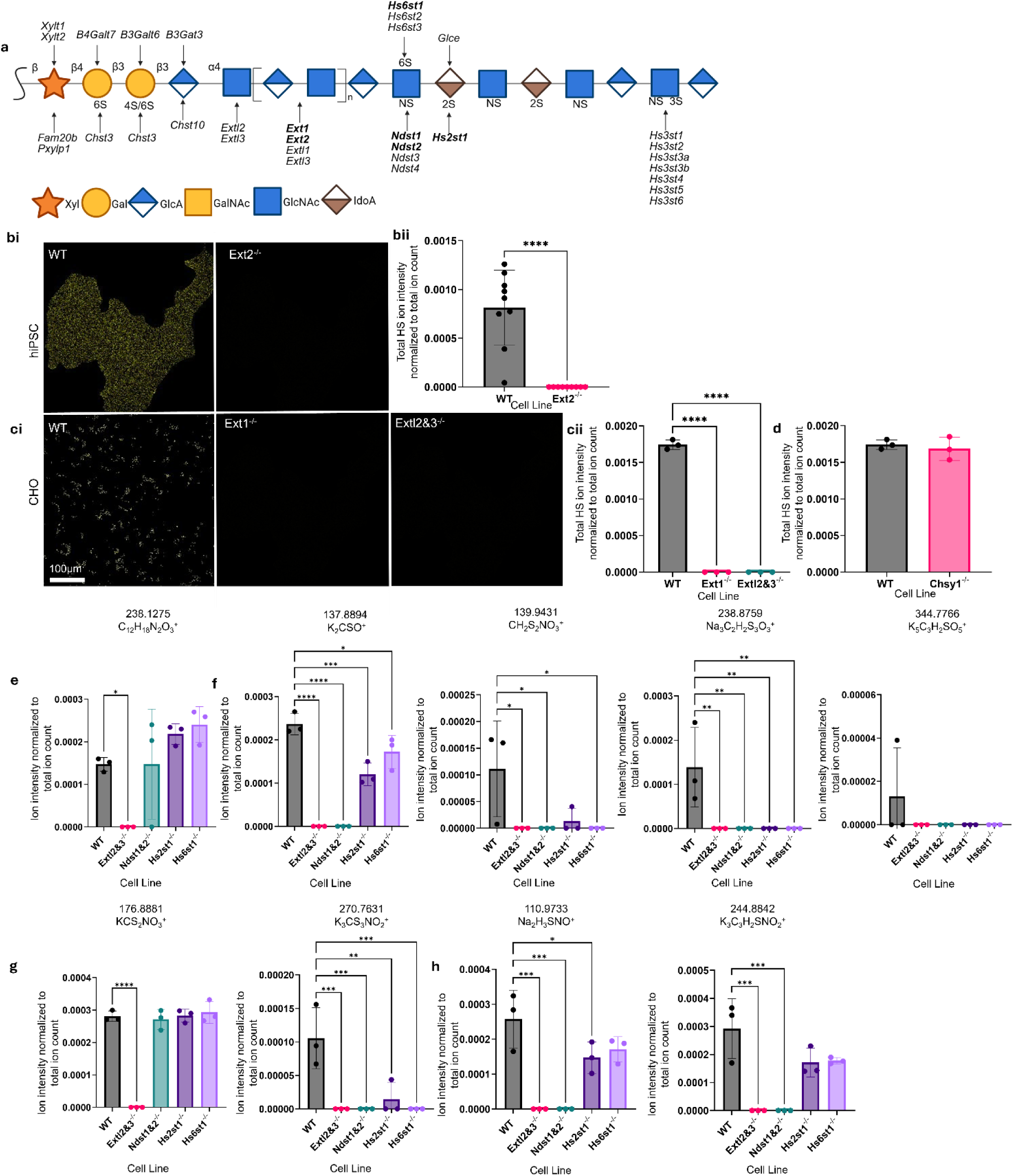
HS discriminatory ions were identified through absence in multiple HS knockout lines (a) HS biosynthetic pathway with manipulated enzymes in bold (bi) Images of summed total HS ion intensity in WT hiPSCs and *Ext2^-/-^* hiPSCs (bii) Comparison of summed HS ion intensity between WT and *Ext2^-/-^* hiPSCs. Absence of HS ions was statistically significant (n=3 of N=3) (unpaired t-test,*p=<0.0001)* (ci) Images of summed HS ion intensity in WT and multiple HS knockout CHO lines (cii) Comparison of summed HS ion intensity between CHO lines. Absence of HS ions was statistically significant in both HS knockout CHO lines (n=3) (one way ANOVA with multiple comparisons, *p=<0.0001* for both *Ext1^-/-^* and *Extl2&3^-/-^*) (d) Comparison of total HS ion intensity between WT and CS knockout CHOs showed no significant difference (n=3) (e-h). Comparison of individual HS ion intensity in Extl2&3^-/-^ (complete HS knockout), *Ndst1&2^-/^*^-^ (loss of *N-*sulphation), *Hs2st1^-/-^* (loss of *2-O-*sulphation) and *Hs6st1^-/-^* (loss of *6-O-*sulphation) to enable relation of ion identity to structure (N=3). (e) Non-sulphated ions identified through lack of sulphate in assignment and absence only in complete HS knockout (f) Ions associated generally with HS sulphation through absence in multiple sulphation modification knockouts (g) Ions associated with sulphate clusters based on presence of a single carbon but multiple sulphates in assignment (h) Ions associated with *N-*sulphation through absence only in *Ndst1&2^-/-^* modification line (N=3).

Here we show the development of analytical pipelines for the integration of ToF and Orbi-SIMS spectra and the first report of genetically modified cell lines for the validation of biomolecule discriminatory ions in SIMS, with previous methods utilizing heavy metals or isotopic labelling^27,28^. The ability to do so for GAGs suggests that the methodology may be applied in the instance of other biological molecules where suitable reference samples are available.

### Assignment of HS ions to specific sulphation patterns

To enable deduction of composition, HS discriminatory ions were assigned to specific GAG structures by comparing ion intensity between WT cells and a library of cells mutant for components of the GAG biosynthetic pathway (Figure 3a)^23^. For clarity, those cells mutant for modification, rather than polymerisation enzymes are defined as specific modification mutants.

A single ion, 238.1275 m/z putatively assigned to C_12_H_18_N_2_O_3_^+^, was found to be associated with the unmodified HS backbone due to (i) its absence only in complete HS knockout lines and (ii) presence across modification specific mutants and (iii) the lack of sulphate in its assignment (Figure 3e).

Four ions (137.8894 m/z K_2_CSO^+^, 139.9431 m/z CH_2_S_2_NO_3_^+^, 238.8759 m/z Na_2_C_2_H_2_S_2_O_2_^+^ and 344.7766 m/z K_2_C_2_H_2_SO_3_^+^) all with sulphate in their assignment, were absent or significantly reduced in multiple modification mutant lines. Suggesting they are likely derived from multiple sulphate patterns in HS (Figure 3f).

176.9991m/z KCS_2_NO_3_^+^ and 270.7631 m/z K_3_CS_3_NO_2_^+^ contain multiple sulphate species, a single carbon, and could not be linked to a specific structure based on intensity in any mutant lines (Figure 3g). It is hypothesized that the positive salt within the assignment could be interacting with the multiple negatively charged sulphates in the highly sulphated HS to form sulphated clusters, and that the presence of multiple sulphate species is the result of inter- and intra-molecular bonds. This has previously been observed in SIMS and is likely a method specific artefact^29^.

Finally, two ions (110.9733 m/z, Na_2_H_3_SNO^+^ and 244.8842 m/z K_3_C_3_H_2_SNO_2_^+^) were identified which could be linked specially to *N-*sulphation, being (i) present in HS and heparin standards, (ii) absent from CS/DS and HA standards, (iii) absent in cells lacking HS and those lacking N-sulphation enzymes *Ndst1^-/-^* / *Ndst2^-/-^*) (Figure 3h).

### Validation of a defined set of ions to identify CS in complex biological samples

A list of 10 CS discriminatory ions was generated based on (i) presence in CS reference spectra (ii) detection in WT cells and (iii) absence in all biological replicates of all cell lines lacking CS synthesis (supplementary figure 5). As with HS, validation of these ions utilised a series of previously characterised CHO lines with modified or absent CS biosynthesis^23^ (Figure 4a). This was initially completed through comparison of WT and complete CS knockout CHO cells. At the time of writing, no CS knockout hiPSC line was available. In WT cells, total CS ion intensity appeared uniform across the cell surface (figure 4bi). The loss of CS ion intensity in *Chsy1^-/-^* CHOs was statistically significant (*p=0.0183* (figure 4bii).

**Figure 4:**
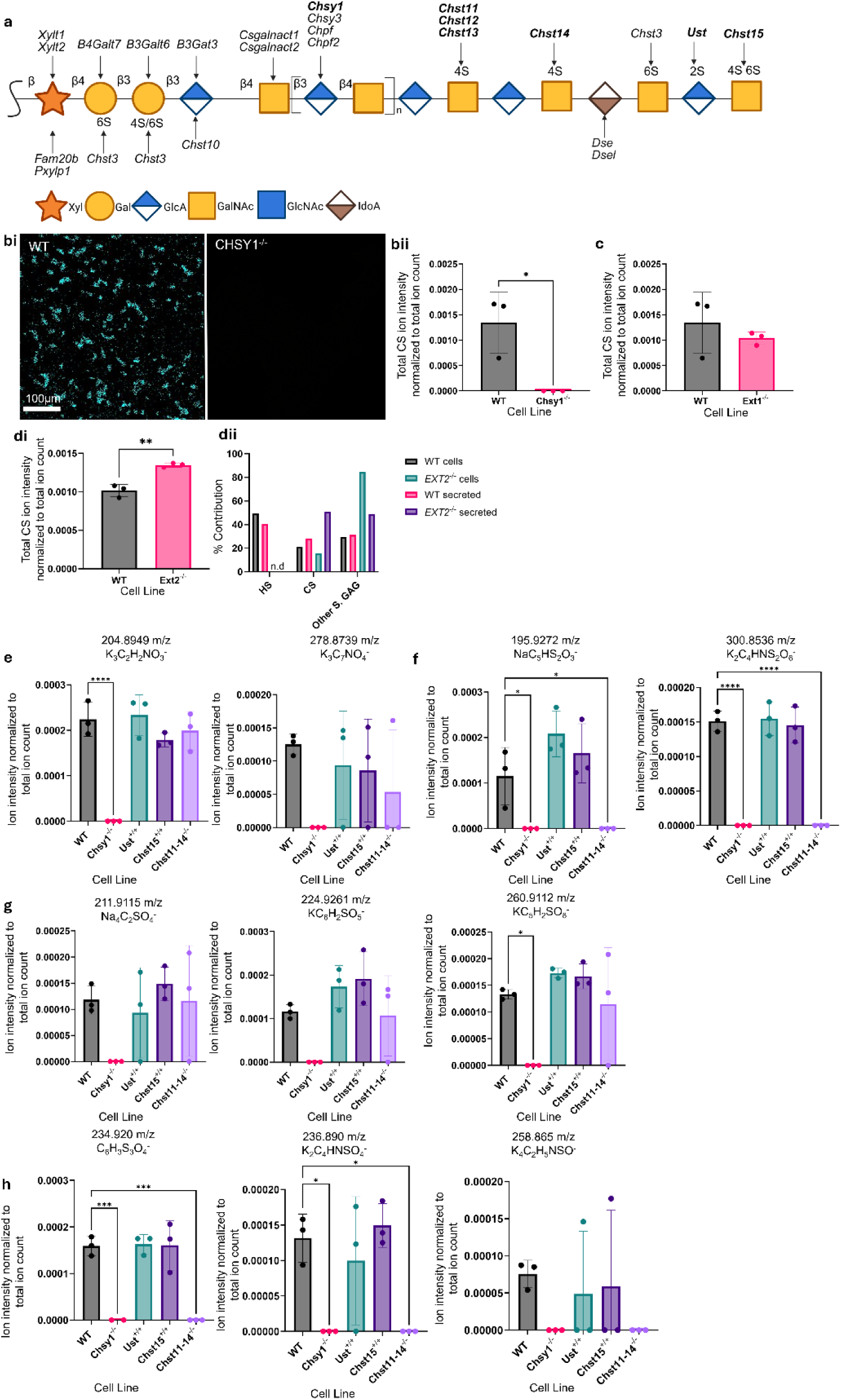
CS discriminatory ions were identified through absence in multiple CS knockout lines (a) Schematic of CS biosynthetic pathway. Modified enzymes are highlighted in bold (bi) Summed CS ion image for WT (left) and CS deficient *Chsy1^-/-^* (right) CHO cells (bii) Comparison of total CS ion intensity between WT and CS deficient CHO cells. Absence of CS ions was significant compared to WT (*p=0.0183,* unpaired t-test) (n=3) (c) Comparison of total CS ion intensity between WT and HS deficient CHO cells (N=3). No significant difference was observed (di) Comparison of total CS ion intensity between WT and HS deficient hiPSCs. CS ion intensity was significantly increased in *Ext2^-/-^* hiPSCs (*p=0.0026,* unpaired t-test) (n=3) (dii) Metabolic radiolabel (Na_2_^35^SO_4_), GAG extraction, differential digestion and size-exclusion chromatography assessed HS, CS and other sulphated GAG (S.GAG) in WT and *Ext2^-/-^* hiPSCs identifying an increase in secreted CS and other sulphated GAG in *Ext2^-/-^* hiPSCs (see also supplemental Figure 3). (e-h) Comparison of the normalized ion intensity for CS ions from wildtype cells, knockouts of CS (*CHSY1^-/-^*) and *4-O-*sulphation (*Chst11-14^-/-^*), and knock-in of 2-4*-O*-sulphation (*Ust*) and 4-6*-O-*sulphation (*Chst15*) (N=3) (e) Non-sulphated ions absent only in cell lines associated with an overall loss or reduction of CS (f) Di-sulphated ions absent in 4-*O-*sulphation knockout (g) Single sulphated species which could not be assigned to a specific sulphation motif (h) Ions absent in 4-*O-*sulphation knockout only.

To further validate the discriminatory nature of these CS ions *in-situ*, their intensity was evaluated in cells lacking HS. No significant difference in total CS ion intensity was observed in HS deficient CHO lines (figure 4c). In *Ext2^-/-^* hiPSCs, a significant increase in total CS ion intensity was observed (figure 4di), consistent with the increased levels of chondroitinase ABC-sensitive material observed following radiolabelled GAG analysis (figure 4dii).

### Assignment of CS discriminatory ions to specific sulphation patterns

A comparison of proposed individual CS discriminatory ion intensity was completed between WT and previously characterised CS modification mutant CHO lines. This included knockout of *Chst11-14* to remove the 4-*O* sulphation motif within CS, knock-in of *Ust* to increase 2,4-*O*-di-sulphated disaccharides or *Chst15* to increase 4,6,-*O*-di-sulphated disaccharides ^23^ with the aim of identifying sulphation motif specific CS ions.

Both 204.8949 m/z, K_3_C_2_H_2_NO_3_^-^ and 278.8739 m/z K_3_C_7_NO_4_^-^ were (i) present in CS reference spectra (ii) absent in a complete CS knockout and (iii) showed no significant intensity changes in modification specific mutants (Figure 4e). In accordance with their non-sulphated putative assignments, this suggests that both ions are associated with unmodified CS disaccharide structures. The presence of nitrogen in both assignments suggests association with GalNAc resides.

195.9272 m/z NaC_5_HS_2_O_3_^-^ and 300.8536 m/z were (i) significantly reduced in the combined *Chst11-14^-/-^* line but (ii) had multiple sulphates in their assignment (Figure 4f). This suggests a potential association with a 4-*O*-sulphated motif and either a second sulphated motif which could not be identified using the currently utilised lines, or sulphate clusters within the GAG chain.

211.9115 m/z Na_4_C_2_SO_4_^-^, 224.9261 m/z KC_6_H_2_SO_5_^-^, 260.9112 m/z KC_5_H_2_SO_8_^-^ (i) showed no significant difference in ion intensity in any modification mutant compared to WT and (ii) had a single sulphate in their assignment (Figure 4g). This indicates that, as in HS, multiple ions have been identified that are liberated from several different sulphated structures in CS.

A final group of ions, containing 234.920 m/z C_6_H_3_S_3_O_4_^-^, 236.890 m/z K_2_C_4_HNSO_4_^-^, and 258.865 m/z K_4_C_3_H_5_NSO^-^ were (i) absent only in only the *Chst11-14^-/-^* modification mutant (ii) contain sulphate in their assignment, supporting their association with a 4-*O*-sulphated motif (Figure 4h).

### SIMS enables concurrent spatial and compositional analysis in tissue

We applied the discriminatory ion sets described above to SIMS data obtained from individual glomeruli of a previously characterized mouse model of streptozotocin (STZ) induced type 1 diabetes^30^ to demonstrate the transformative value of the ion sets for spatially resolved analysis of GAGs.

ToF-SIMS analysis was completed on archive samples (FFPE blocks, sectioned and mounted). Spectra were extracted from a region of interest associated with a single glomerulus from three healthy and three diabetic mice (Figure 5a). In this case, both the composition and spatial arrangement of the GAGs within the glycocalyx underpin their biological role, thus, applying SIMS to study GAGs is crucial in developing the understanding of GAGs in diabetes. Importantly, the workflows developed here are not disease specific and could be explored in other tissue types and disease states to improve the link between spatial localisation and GAG function including, but not limited to, the vascular wall. Furthermore, the speed of SIMS could be exploited for tissue analysis to enable high throughput biomarker or therapeutic target discovery and the monitoring of treatment efficacy, with the presented cellular images taking approximately 90 seconds to acquire and higher resolution glomerular images taking approximately 30 minutes. Finally, the low pixel size (244 nm) enabled region of interests to be selected corresponding to individual glomeruli (diameter » 60 mm), demonstrating the high spatial precision capabilities of the technique.

**Figure 5.**
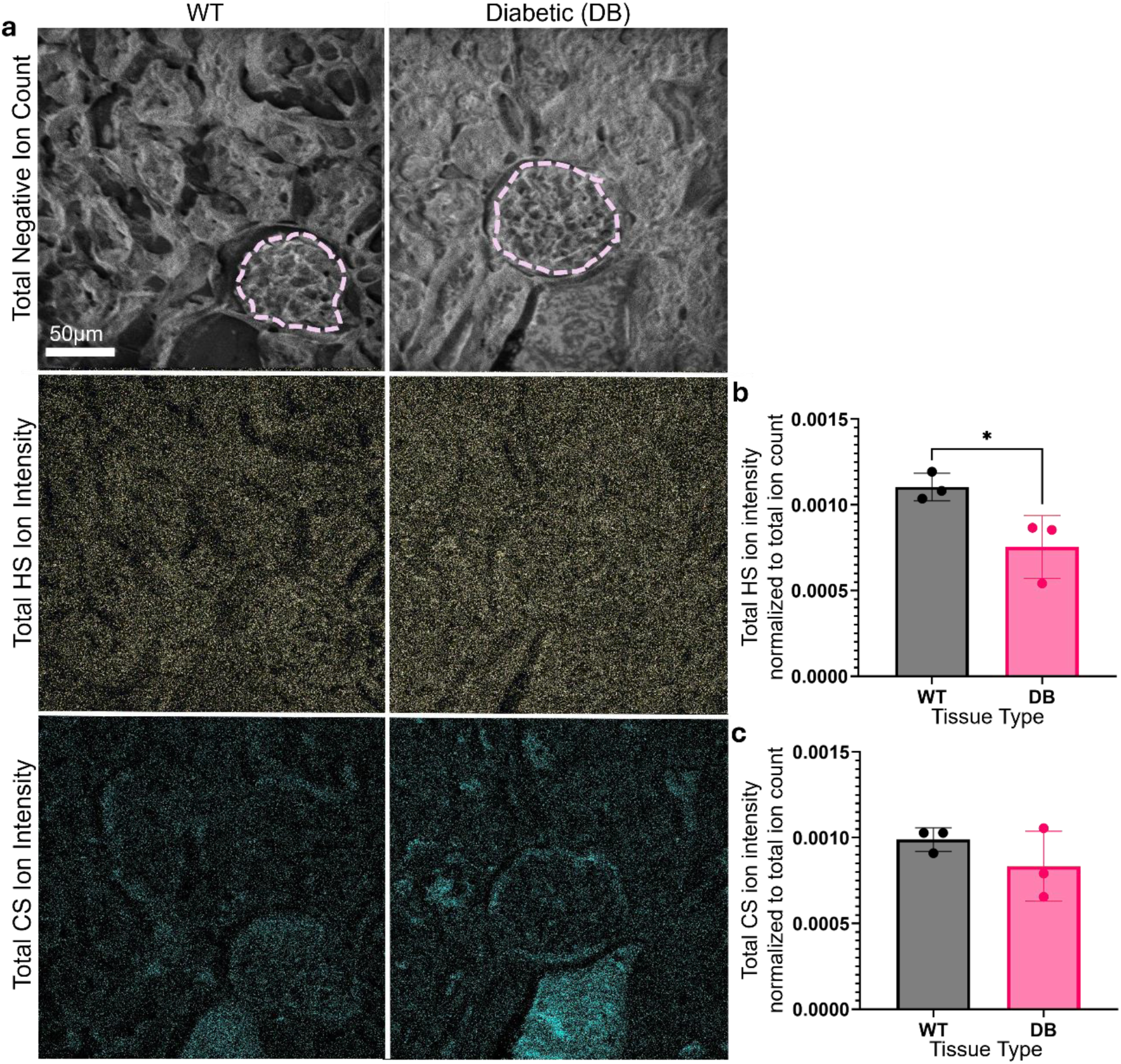
ToF-SIMS can identify GAG discriminatory ions in FFPE tissue sections (a) Total negative ion count, total HS and total HS spatial images of individual glomeruli (ROI outlined in pink dashed line) in WT and diabetic (DB) mice (b) Comparison of HS ion intensity in glomeruli (intensity taken from ROI). Total HS ion intensity was significantly reduced in DB compared to WT glomeruli (*p=0.0394,* unpaired t-test) (c) Comparison of total CS ion intensity in glomeruli (intensity taken from ROI). No significant difference in CS ion intensity was observed (N=3).

Of the previously identified ions, all HS ions except 270.76 and 344.78 m/z could be identified in control and diabetic tissue. Total HS ion intensity was significantly reduced (*p=0.0394*) in diabetic glomeruli compared to control (figure 5b). This is in agreement with previous literature describing a heparinase induced loss of HS from the glomerular basement membrane in an STZ-induced diabetic murine model ^31,32^. All CS ions could be found in both control and diabetic glomeruli and total CS ion intensity showed no significant difference between groups (figure 5c). Whilst we have previously been able to confidently identify ions that discriminated between purified CS and DS^7,20^ and shown in cellular systems both potential CS and DS discriminatory ions and labelled them accordingly (supplementary table 5 & 6). We recognise that given the essential difference between CS and DS is the presence of IdoA in DS (with its epimer, GlcA in CS), that in complex biological samples these ions have the potential to originate from either. We are continuing to investigate how discriminatory ions are liberated from such similar structure.

## Discussion

Previous application of SIMS for *in-situ* GAG analysis is limited and has typically centred around the use of sulphate species that are not unique to GAGs ^21,22^. However, our lab has previously shown that SIMS can discriminate between picogram quantities of individual GAG standards, suggesting that the SIMS ions produced from GAGs are informative of GAG type and composition ^7,20^.

In the present study, GAG-derived ions were identified from SIMS spectra associated with 2D-cell culture samples and tissue samples using both purified reference materials and knockout/knock in cell lines. This included ions associated with GAGs generally, individual GAG types, and specific sulphation motifs. These ions were detected in both 2D cell culture samples, healthy and diabetic renal tissue, enabling concurrent spatial and compositional GAG analysis at sub-micron resolution. SIMS has been successfully applied to analyse other biological molecules in a wide variety of tissue types^33–35^, suggesting that SIMS analysis of GAGs could also be translated to diverse clinically relevant samples. The rapid analysis times (as little as 90 seconds for the presented cellular images) make SIMS analysis ideal for high throughput biomarker discovery projects and for exploring the fundamental biological roles of GAGs. Significantly, this method avoids the use of labels or digestion for GAG detection, release or characterisation and is therefore unbiased. Enzymatic digestion and incorporation of labels can be hard to control and typically have selectivity within the structural diversity of GAGs^23,36^.

Unlike previous validation of SIMS ions which have utilized heavy metal or isotopic labels^27,28,37^, we have taken a genetic approach to ion validation, confirming ions of interest are absent in complex biological systems with known loss or gain of GAGs or GAG motifs. As with the described data integration process, the identification of GAG discriminatory ions in this way serves as an example of a principle that could be applied to any biomolecule known to ionize in SIMS, to aid in the identification of discriminatory or associated ions within complex and convoluted spectra.

Whilst the described library shows, for the first time, that SIMS can identify GAG specific ions *in-situ*, it does not contain a fully comprehensive library of validated ions for all GAG types of sulphation motif. Development of additional reference samples, including synthetic GAG oligosaccharides or genetically modified materials from different tissues or disease states, will enable expansion of the ion library through the identification of additional SIMS ions that are informative of different GAG compositions.

## Methods

### Preparation of individual GAG standards

Sodium adduct individual GAG standards were reconstituted to a concentration of 5mg/ml in ultrapure water (HS from porcine mucosa, Iduron; DS from porcine mucosa, Iduron; CS from porcine mucosa (Sigma-Aldrich), KS from bovine mucosa; University of Uppsala (gift from Prof. Ulf Lindahl); HA from *Streptococcus Equi,* Sigma-Aldrich). As described in Hook et al., 2021, for each sample, 1 µl of solution was deposited onto a poly-_L_-lysine coated slide (Poly-prep, Sigma-Aldrich) and allowed to dry at room temperature. After drying, a further 1 µl of solution was deposited onto the dried droplet. This was repeated a third time^20^.

### Orbi-SIMS of individual GAG standards

The Orbitrap analyser was calibrated using a silver sample according to a previously described method^14^. Orbi-SIMS analysis of the GAG standards was conducted as previously described^7^.A 30 keV Bi_3_^+^ liquid metal ion gun was used as the primary ion source. Data was collected using the Orbitrap™ analyser over an analysis area of 200 × 200 μm^2^ for 30 scans. The cycle time was set to 200 μs. The spectra were collected over a mass range of *m*/*z* 80–1000 in positive and negative polarity. One area per sample was analysed.

### Ion Assignment

Assignments of peaks was completed the series of criteria outlined previously^7^. In short, this limited the assignment to only contain C, S, O, N, Na or H and the number of species to those chemically possible in individual disaccharide units. Based on their shared presence in HA, sulphate was excluded in the assignment of all common GAG peaks. Peaks were assigned within a 5ppm deviation of the Orbi-SIMS reference spectra.

### Cell preparation for SIMS

For hiPSC culture, 13mm polystyrene tissue culture plastic coverslips (Nunc 150067) were UV sterilised for 30 min. Coverslips were coated with 300 μl 0.5 μl/ml vitronectin. hiPSCs were passaged as described (see supplemental methods). Following re-suspension using manual force, cells were resuspended in 1ml of fresh E8 and 50,000 cells seeded per coverslip. Cells were grown to approximately 70% confluency.

GAG synthetic CHO lines were generated and validated using disaccharide analysis by the Copenhagen Centre for Glycocalyx Research. WT, Extl2&3^-/-^, NDST1&2^-/-^, HS2ST1^-/-^, HS6ST1^-/-^, CHSY1^-/-^, UST^+/+^, CHST15^+/+^ and CHST11-14^-/-^ cells were maintained in suspension culture as detailed previously^23^.

18 mm glass coverslips (Fisherbrand) were sterilised by incubation in 70% ethanol overnight at room temperature before airdrying at room temperature. Cells were passaged using manual pipette force before centrifugation at 300 x g for 2 min. A manual cell count was completed and 50,000 cells were seeded per coverslip. Cells were grown to 70% confluency before fixing with 4% PFA for 20 min at room temperature.

### Optimization of sample preparation

Methods of sample preparation for SIMS of GAGs were optimized on WT hiPSCs. Cells were seeded and grown as described above.

Untreated (UT) cells had media removed and coverslips were allowed to airdry at room temperature. Ammonium formate (AF) washing was completed in accordance with previously used methods of sample preparation for SIMS of biological samples^38^. Cell culture media was aspirated and cells washed for 3 x 1 min with 150 mM AF and allowed to try at room temperature. Chemical fixation (FX) was used to determine if SIMS of GAGs is possible on fixed tissue, in accordance with preparation for other imaging modalities such as fluorescence microscopy. This was completed by aspiration of cell culture media, followed by 4% PFA fixation for 20 min at room temperature and 3 washes with PBS. Freeze drying (FD) is another standard method of sample preparation for SIMS of biologicals. Cell culture media was aspirated. Samples were stored at -80 °C for 12 hours, followed by freeze drying for 12 hours. Combinatorial approaches were also explored, including FX followed by FD (FXFD) and AF, followed by FX and FD (AFFXFD). Three technical replicates per condition were used for optimisation of sample preparation.

### ToF-SIMS of cellular samples

Data was collected using a ToF-SIMS 5 (IonToF GmbH, Germany) run in high current bunch mode. A 30 keV Bi_3_^+^ liquid metal ion gun was used as the primary ion source. The main chamber was maintained at a pressure of 5 x 10^-8^ mbar. A low energy electron flood gun was used for charge compensation on the sample surface. A mass range of m/z 0-899 was collected in both positive and negative polarity. Ions were predominantly present below m/z 400. A cycle time of 100 µs was used. For all cell lines, 3 x 500 µm x 500 µm regions were imaged at a pixel density of 256 x 256 using the random raster mode. 1 frame per patch was collected in 15 scans per measurement. Analysis of each region took approximately 60 s to acquire the spectra and associated images.

### Tissue preparation for SIMS

All animal experiments were approved by the UK Home Office. An STZ induced model of type 1 diabetes in DBA2/J mice was generated as previously described^30^. In brief, STZ was given i.p at 50mg/kg for 5 consecutive days. Mice were killed at 8 or 9 weeks post-STC injections. Mice with glycemia of ≥15mmol/l were considered diabetic. 4µm thick sections of formalin fixed, paraffin embedded kidneys from three control and three diabetic mice were mounted onto superfrost slides. Samples were deparaffinated using the following wash steps: 2 x 10 min in xylene, 5 min in 100%, 95% and 70% ethanol, 3 min in PBS and 3 min in distilled water. All slides were dried overnight at room temperature prior to SIMS analysis.

### ToF-SIMS of individual glomeruli

Data was collected using a ToF-SIMS 5 (IonToF GmbH, Germany) run in delayed extraction mode. Individual glomeruli were identified in tissue sections using the brightfield camera present in the instrument. A 30 keV Bi_3_^+^ liquid metal ion gun was used as the primary ion source. The main chamber was maintained at a pressure of 5 x 10^-8^ mbar. A mass range of m/z 0-899 was collected in both positive and negative polarity. A cycle time of 100 µs and a delayed extraction time of -0.005µs was used. A 250 µm x 250 µm region at a pixel density of 1024 x 1024 was imaged using the random raster mode. 20 frames per patch were collected in 1 scan.

### Spectral Analysis

Surface Lab version 6.8 (IonToF GmbH) was used for the analysis of all spectra. The region of interest tool was used to identify cellular regions in the total negative ion count image. Where possible this was around cell clusters. However, if cells were sparse, individual cells were selected, and a combined region of interest was generated. Positive spectra were calibrated to CH_3_^+^, C_2_H_5_^+^ and C_3_H_8_^+^ and negative spectra to C_2_^-^, C_3_^-^ and C_4_H^-^. All ion intensity values were normalized to total ion count.

### Identification of GAG discriminatory ions

A peak search of a single replicate of UT, WT hiPSCs was completed for ions in a mass range of 75-400 m/z with a minimum count of 1000 to create an initial peak list. Corresponding ToF-SIMS peaks from different cellular samples were identified within ±100 ppm deviation from a peak’s central mass. To confirm these were GAG related, equivalent Orbi-SIMS peaks from the GAG reference spectra were identified based on their central mass being within ±5 ppm of the mass value of the ToF-SIMS cellular peak.

This process was repeated for all individual GAG standards to identify peaks present in both the standards and WT hiPSCs. Common GAG ions were those present in the reference spectrum of at least one sulphated GAG and hyaluronic acid and identification in at least n=3 WT hiPSC spectra. Individual GAG discriminatory ions for all GAG types were identified based on their presence a single standard reference spectra (absent in the spectra of other GAG types) and detection in n=3 of WT hiPSCs.

To validate all potential individual GAG ions, their presence was confirmed in at least two biological replicates of WT cells and their absence confirmed in all biological replicates of corresponding knockout cell lines. Absence was defined as an undetectable peak or an area of less than 500 counts. HS discriminatory ions were validated through absence in Ext2^-/-^ hiPSCs and Ext1^-/-^ CHOs and CS discriminatory ions were confirmed through absence in CHSY1^-/-^ CHO cells.

To enable relation of GAG ion identify to more specific saccharide structures, the absence of ions was assessed in discrete knockout CHO lines. For HS, NDST1^-/-^ cells were used to assess ions with a relationship to *N-*sulphation. HS2ST^-/-^ and HS6ST2^-/-^ were used to determine if HS ions could be linked to *2-O* or *6-O* sulphation respectively. For CS, CHST11-14^-/-^ cells were used to relate ion identity to *4-O* sulphation, UST^+/+^ to *2-O* sulphation in CS/DS and CHST15^+/+^ cells to identify *6-O* sulphation^23^.

### Image Analysis

All images were exported from Surface Lab 6.8 software as an Ascii text file. A Fiji macro was written (Supplementary information) to convert the text file to a .tiff image file for display purposes. The region of interest tool was used to select cellular regions in the total negative ion count image and applied to all images. The sum tool was used to combine all images of interest into a stack per polarity. Images were normalized to total ion count using the image calculator tool to divide the image of interest by the corresponding polarities total ion count image. Look up tool was used to apply colour to each image.

### Statistical Analysis

Prism version 10.3.0 was used to plot all graphs and perform statistical analysis. Data points correspond to individual replicates ± standard deviation. Further description of if this refers to technical or biological replicates is outlined in figures. All statistical tests used are described in figures.

## Acknowledgements

The authors would like to thank the SIMS facility at the University of Nottingham for access to instrumentation. Figures created in BioRender. Merry, C. (2026) https://BioRender.com/omfv1f8, https://BioRender.com/ob607nd, https://BioRender.com/u9dj76n. Funding for this work was provided by the Biotechnology and Biological Sciences Research Council (BBSRC) Impact Acceleration Account administrated through the University of Nottingham (CLRM/ALH), the Engineering and Physical Sciences (EPSRC) Next Generation Biomaterials Discovery Award (EP/N006615/1; CLRM, JLT), the BBSRC Strategic Longer and Larger award, GlycoWeb (BB/Y00311X/1; CLRM/AH/KPA), an award from Kidney Research UK (LM/CLRM/ALH/KPA) and the EPSRC (EP/Y002423/1; LKM/ALH). PhD doctoral training awards were received from the Biotechnology and Biological Sciences Research Council (LKM/KPA/CLRM/ALH).

## Supplementary Information

### Supplementary Methods

#### hiPSC lines origin, characterization and maintenance

REBL-PAT human induced pluripotent stem cells (hiPSCs)^39^ cells were a generous gift from Prof. Chris Denning, Biodiscovery Institute, University of Nottingham. The EXT2^-/−^ hiPSC line was generated from the REBL-PAT-hiPSC line through nucleofector electroporation (Lonza 4D, program CA-137) of 1 μg total DNA and homologous recombination followed by antibiotic selection (Figure S2)^40^. The EXT2^-/−^ hiPSC line was further validated through PCR and sanger sequencing (Figure S2). All hiPSC lines were seeded on vitronectin recombinant human protein and cultured in E8 medium in a humidified atmosphere (5% CO_2_) at 37°C. Cells were passaged twice a week using TrylpLE Select (1X) and E8 supplemented with 10 μM Y-27632 Rho kinase inhibitor (Stem Cell Technologies). Human stem cells were subject to routine pluripotency testing using BD Stemflow Human and Mouse Pluripotent Stem Cell Analysis Kit (BD Biosciences) as recommended by the manufacturers, or by immunostaining against OCT3/4, SOX2 and NANOG using the standard immunostaining protocol as detailed below. Karyotype stability was also monitored (Figure S2).

#### Immunocytochemistry

REBL-PAT (wild type, WT) and EXT2^-/−^ hiPSCs were cultured on recombinant vitronectin coated tissue culture plastic, fixed with 4% paraformaldehyde, with staining performed essentially as described previously^41^ (and see table S1 For antibodies used). Where required, slides were treated with 2 mIU heparinase I, II, and III (Iduron) in PBS with Ca^2+^ and Mg^2+^ (Gibco) for 1 h at room temperature to digest HS prior to permeabilization. Cells were imaged using a high-content Microscope (Operetta, Perkin Elmer).

#### Flow Cytometry

Flow cytometic detection of GAGs was carried out essentially as described^42^. Cells were treated with gentle cell dissociation reagent for 10 min at 37°C and washed in 1X DPBS. Samples were pelleted at 160 x g for 4 min and resuspended in fluorescence-activated (FACS) buffer (1% sterile filtered bovine serum albumin in 1X DPBS). 40,000 cells per staining condition were transfered into a V bottom 96 well plate. hiPSCs were incubated with primary antibody (Supplementary Table 1) diluted in FACS buffer for one hour at 4 °C followed by three FACS buffer washes. Samples were then incubated with secondary antibody (AlexaFluor 488-tagged goat anti-mouse IgG, anti-mouse IgM) diluted 1:500 in FACS buffer for one hour at 4°C, followed by three FACS buffer washes. All FACS was completed on a FC500 Flow Cytometer (Beckman-Coulter) and data analysis completed on Kaluza.

#### 2-Aminoacridone (AMAC) Labelling for High Performance Liquid Chromatography Disaccharide Analysis

Preparation and analysis was essentially carried out as described previously^43^. hiPSCs were seeded at 20,000 cells/cm^2^ on vitronectin coated plates and cultured in E8 media for 72 hours. Conditioned media was collected and pooled from both 48 and 72 hours. At 72 hours, cells were washed with 1X DPBS and lysed using 1% (vol/vol) Triton X-100 for 2 hours at room temperature on an orbital shaker. Media and cell extracts were treated with 100 µg/ml Pronase (Roche Diagnostics, 0165921001) for 4 hours at 37°C. Samples were then incubated for 15 min at 90°C to deactivate pronase. Cells were incubated with 14 µg/ml DNAse1 (Sigma, Dn25) in 10 mM MgCl_2_ (Sigma, M8266) overnight at 37°C to prevent DNA contamination. Preparations were loaded onto DEAE-Sephacel columns (Sigma, I6505). 1.5 M NaCl, 20 mM NaH_2_PO_4_.H_2_O at pH 7.0 was used to equilibrate columns. Columns were washed with 50 x column volume of 0.25 M NaCl, 20 mM NaH_2_PO_4_.H_2_O pH 7.0 to remove HA. Sulphated GAGs were eluted using a 5 x column volume of 1.5M NaCl, 20 mM NaH_2_PO_4_.H_2_O pH 7.0 and desalted using PD-10 Sephadex G-25M pre-packed columns (GE Healthcare, 17-0851-01). HS chains were digested using 0.8 mIU of heparinase I, II and III (Iduron) in 100 µl of 0.1 M sodium acetate and 0.1 mM calcium acetate pH 8.0. CS and DS chains were digested using 0.8 mIU chondroitinase ABC (AMSBio AMS.E1208-02) diluted in 100 µl 50 mM Tris buffer, 50 mM sodium chloride pH 7.9. Cells were incubated with GAG lyases for 12 hours at room temperature. Resulting disaccharides were freeze-dried, resuspended in 10 µl of 0.1 M 2-aminoacridone (Sigma, 06227) and incubated at room temperature for twenty minutes. Samples were incubated with 10 µl of 1M NaBH_3_CN overnight at room temperature. AMAC-labelled samples were run in duplicate and separated using a Zorbax Eclipse XDB-C18 RP-HPLC Column (35 µM, 2.1 mm x 150 mm) (Agilent Technologies). Disaccharides were identified and quantified based on comparison to AMAC-labelled disaccharide standards (Iduron)^36^.

#### Metabolic (Na_2_ ^35^SO_4_) Radiolabelling of GAGs

Metabolic incorporation of radiolabel into GAGs followed by extraction and characterisation was used to verify the lack of HS in the EXT2^-/−^ hiPSC line. REBL-PAT (WT) and EXT2^-/−^ hiPSC lines in E8 media on vitronectin were incubated with 300 μCi Na_2_[^35^S]sulfate (PerkinElmer) in E8. After 18 h, ^35^S-labeled proteoglycans were purified from cell extracts by DEAE ion-exchange chromatography and gel chromatography on Superose 6 columns as described previously^44^. Size distribution of [^35^S]sulfate-labeled, NaOH-released GAGs was analyzed by gel chromatography on Superose 6 columns in 1% Triton X-100, 1 m NaCl, 50 mm Tris-HCl, pH 7.5. Fractions of 0.5 ml were collected and analyzed by scintillation counting. The mixed GAG population was separated before and after digestion with either a combination of heparinase I, II and III (Iduron) or chondroitinase ABC (AMS Bio) to discriminate between different GAG populations. The contribution of each component (HS, CS/DS or an un-identified GAG that was resistant to both enzymatic digestions) was calculated for the extracts from both cell types.

---

**Supplementary Figure 1.**
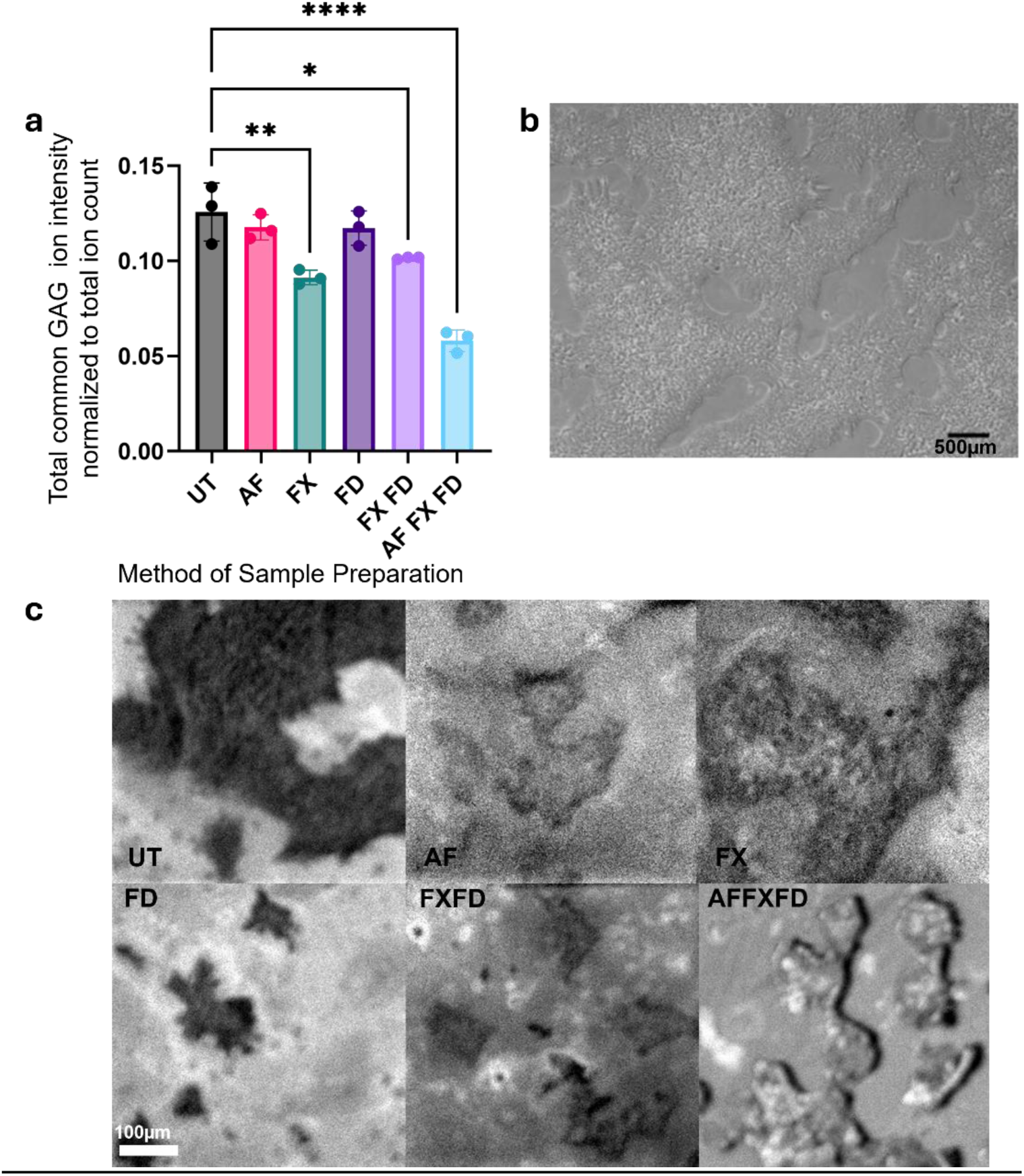
Chemical fixation (4% PFA) is a viable sample preparation method for SIMS of GAGs. Methods of sample preparation (UT: untreated, AF: ammonium formate wash, FX: 4% PFA fixation, FD: freeze dried, FXFD: 4% PFA fixation followed by freeze drying, AFFXFD; ammonium formate wash, 4% PFA fixation followed by freeze drying) (a) Comparison of putative total common GAG ion intensity between sample preparation methods (Ordinary one-way ANOVA with multiple comparisons to untreated samples,n=3) (b) example brightfield image of WT hiPSCs in culture (ci) Total positive ion count of WT hiPSCs under all methods of sample preparation for comparison of morphology to bright field image. UT and FX were most comparable to that of cells in culture.

**Supplementary Figure 2.**
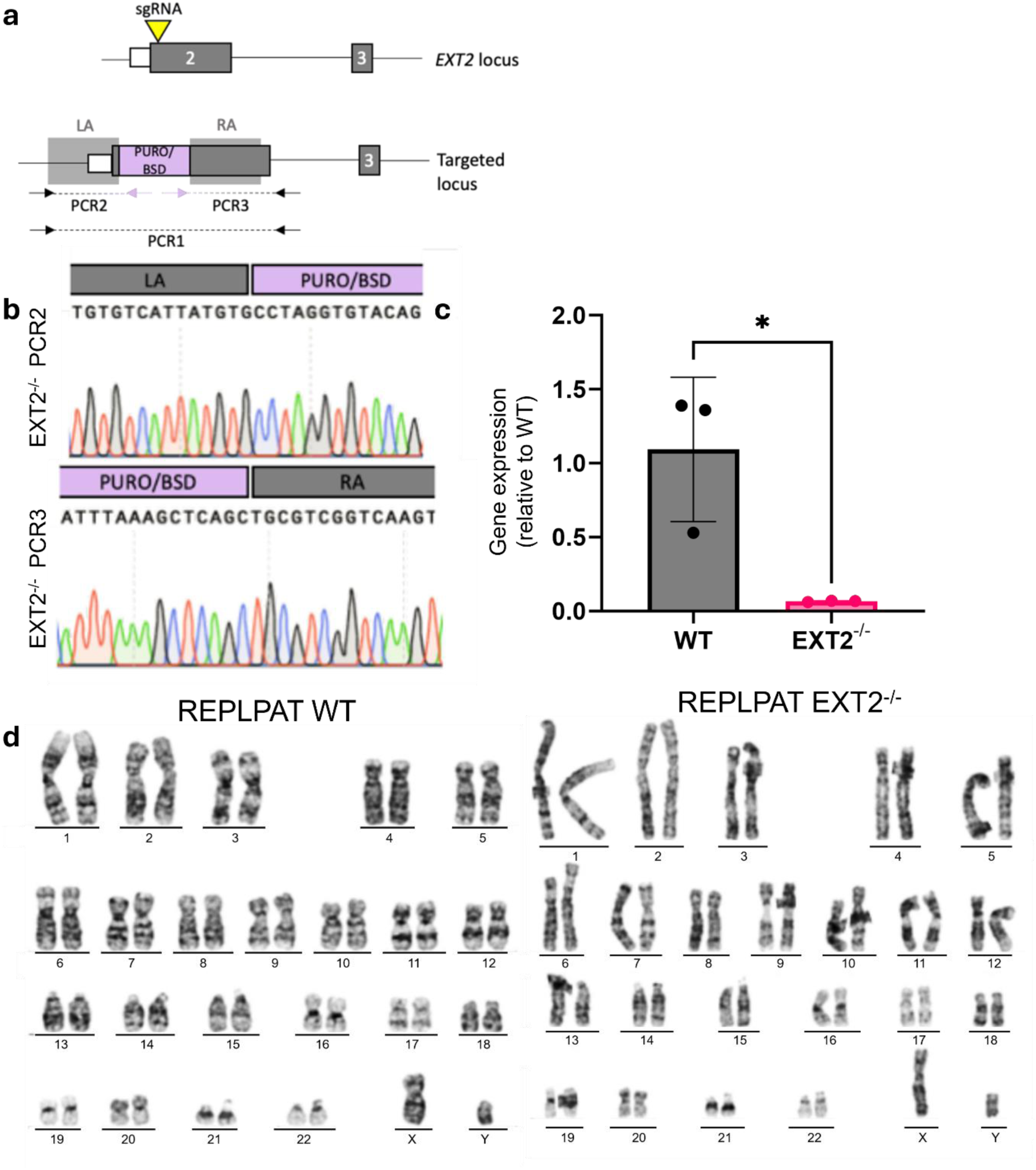
Validation of Ext2^-/-^ hiPSCs (a) Diagram showing insertion of puromycin or blastocidin after the transcription start site of the EXT2 gene. (b) Sanger sequencing of Ext2^-/-^ hiPSC clones using PCR2 and PCR3 amplicons show correct insertion of the antibiotic resistance gene associated with CRISPR-Cas9 targeting strategy (c) qRT-PCR was used to compare EXT2 gene expression between REBL-PAT (WT) and Ext2^-/-^ hiPSCs. EXT2 gene expression was significantly reduced in EXT2 knockout cells. Error bars represent ±SEM from N=3. *p<0.05, Tukey’s multiple comparisons test (d) G-band karyotyping was used to assess genomic integrity in WT hiPSCs (passage 31) and Ext2^-/-^ hiPSCS (passage 25) post targeting. Both cell lines maintained a normal karyotype.

**Supplementary Figure 3.**
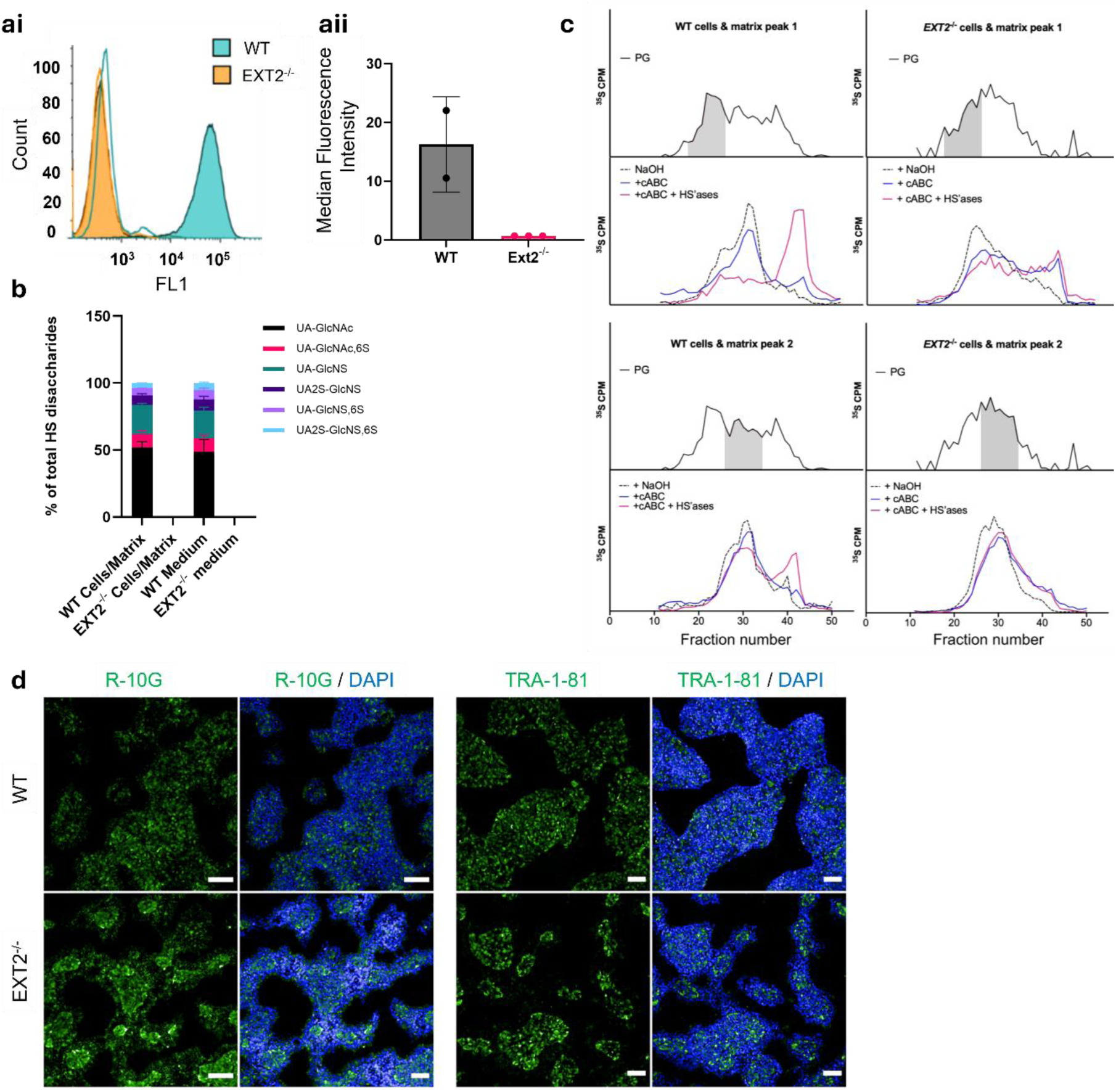
Ext2^-/-^ hiPSCs do not produce HS and display an altered GAG profile. (ai) Flow cytometry was used to assess the display of HS on Ext2^-/-^ hiPSCs. 10E4, which binds a common N-sulfated epitope within HS chains^45^, failed to detect any HS on EXT2^-/-^ cells (orange) in contrast to WT cells (cyan), isotype control in red (aii) Mean fluorescence intensity of 10E4 was normalised to secondary only controls. Error bars represent ± SEM from n=2 (WT) n=3 Ext2^-/-^ from independent experiments (b) HS disaccharide composition of material isolated from WT and Ext2^-/-^ hiPSCs, Material was separately prepared from the cells/matrix and conditioned medium, digested, labelled and fractionated as described previously^44^. Data is mean ±SEM of N=3 biological samples (c) ^35^S-sulphate labelled proteoglycans were isolated from hiPSC cell fractions and conditioned medium by DEAE ion chromatography. Purified proteoglycans from WT (left) and Ext2^-/-^ (right) were analysed by Sepharose 6 size exclusion chromatography. Fractions from peak 1 & 2 (shaded grey) were collected, alkali treated (NaOH) and subsequently digested with chondroitinase ABC and heparinases (alone or in combination). All profiles were normalised to 10,000 CPM (d) Fluorescence imaging of WT and Ext2^-/-^hiPSCs after immunofluorescence staining using R-10G and TRA-1-81 (co-stained with DAPI) to detect KS epitopes. Scale bar = 100 μm.

**Supplementary Figure 4.**
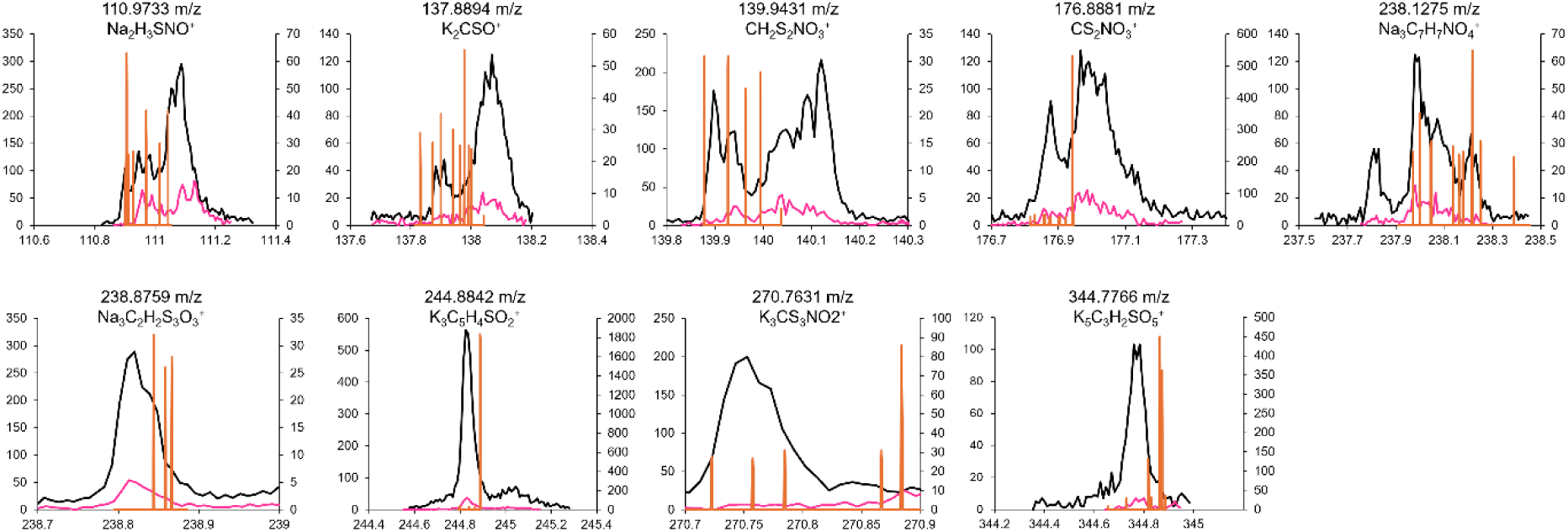
Comparison of counts values of HS reference spectra to WT and Ext2^-/-^ hiPSCs. Overlap of HS discriminatory ions in WT and Ext2^-/-^ hiPSC ToF-SIMS spectra (binned to 10 channels) with HS Orbi-SIMS reference spectra (binned to 1 channel). Secondary axes are used for reference spectra as required (Black – WT cells, Pink – HS KO cells, Orange – HS reference).

**Supplementary Figure 5.**
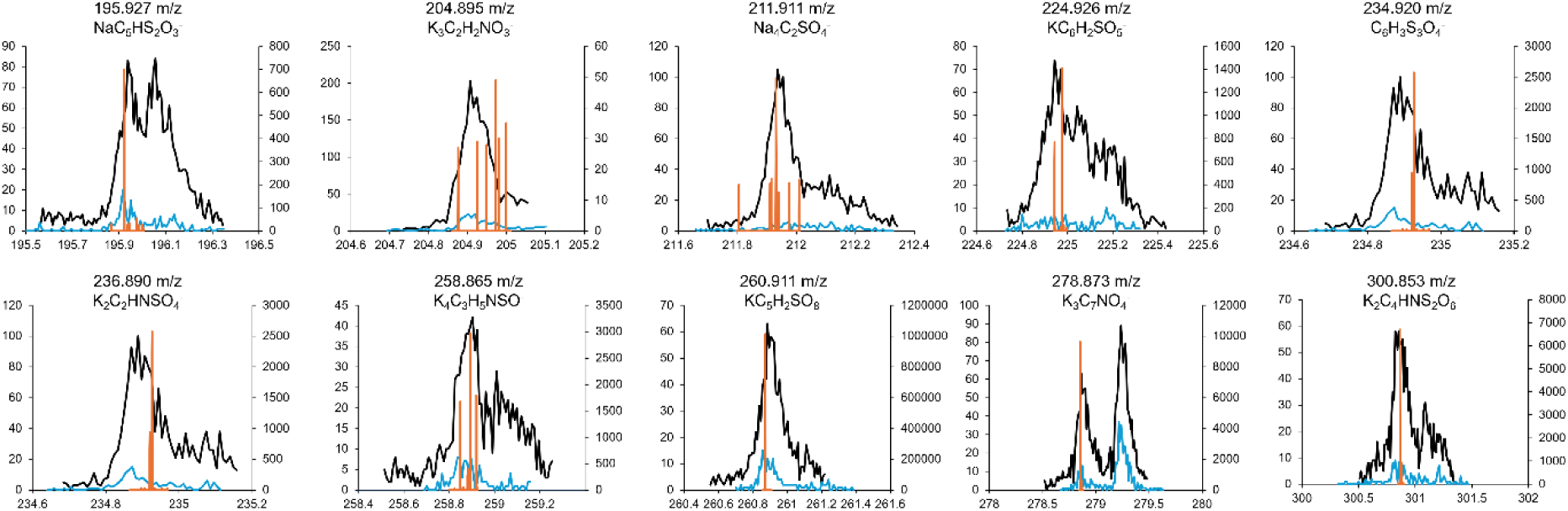
Comparison of CS reference spectra to WT and CHSY^-/-^ CHOs. Overlap of CS discriminatory ions with WT and CHSY1^-/-^ CHO cells ToF-SIMS spectra (binned to 10 channels) with CS Orbi-SIMS reference spectra (binned to 1 channel) Second axis are used for reference peak plotting as required. (Black – WT cells, Blue – CS KO cells, Orange – CS reference).

**Supplementary Table 1.** Antibodies used in this study.

| **Antibody** | **Epitope** | **Supplier** |
| --- | --- | --- |
| 10E4 IgM | Intact N-sulfate HS | AMSBio |
| 3G10 IgG | HS stub remaining after heparinase digestion | AMSBio |
| 5D4 | Sulfated keratan sulfate | Gift from Prof. Ten Feizi, King's College London |
| R-10G | Non/low sulfated keratan sulfate | Merk |
| TRA-1-81 | Keratan sulfated podocalyxin | Thermofisher Scientific |
| Alexa Fluor 488-tagged anti mouse IgG/IgM or anti-rabbit IgG | Secondary antibodies used with the above | Thermofisher Scientific |

**Supplementary Table 2.** Positive common GAG ions identified based on their presence in the reference spectra of hyaluronan, at least one sulphated GAG and WT hiPSCs.

| m/z | Assignment | m/z | Assignment |
| --- | --- | --- | --- |
| 87.1093 | C<sub>2</sub>HNO<sub>3</sub><sup>+</sup> | 164.9173 | Unknown |
| 92.0544 | Na<sub>2</sub>CHO<sub>2</sub><sup>+</sup> | 165.0570 | K<sub>2</sub>C<sub>3</sub>H<sub>5</sub>NO<sub>2</sub><sup>+</sup> |
| 117.9895 | Na<sub>2</sub>C<sub>2</sub>H<sub>2</sub>NO<sub>2</sub><sup>+</sup> | 168.0371 | NaC<sub>5</sub>H<sub>5</sub>O<sub>5</sub><sup>+</sup> |
| 118.9347 | Na<sub>2</sub>C<sub>2</sub>HO<sub>3</sub><sup>+</sup> | 168.9607 | Na<sub>3</sub>C<sub>4</sub>H<sub>4</sub>O<sub>3</sub><sup>+</sup> |
| 119.9280 | Na<sub>2</sub>C<sub>2</sub>H<sub>4</sub>NO<sub>2</sub><sup>+</sup> | 169.9848 | Unknown |
| 121.9221 | Na<sub>2</sub>C<sub>2</sub>H<sub>3</sub>O<sub>3</sub><sup>+</sup> | 170.0561 | NaC<sub>5</sub>H<sub>7</sub>O<sub>5</sub><sup>+</sup> |
| 128.0754 | NaC<sub>3</sub>H<sub>5</sub>O<sub>4</sub><sup>+</sup> | 171.0867 | NaC<sub>4</sub>H<sub>6</sub>NO<sub>5</sub><sup>+</sup> |
| 129.0604 | Na<sub>2</sub>C<sub>4</sub>H<sub>3</sub>O<sub>2</sub><sup>+</sup> | 172.1037 | C<sub>7</sub>H<sub>10</sub>NO<sub>4</sub><sup>+</sup> |
| 130.0657 | Na<sub>2</sub>C<sub>4</sub>H<sub>6</sub>NO<sup>+</sup> | 174.0260 | Na<sub>2</sub>C<sub>5</sub>H<sub>6</sub>NO<sub>3</sub><sup>+</sup> |
| 131.9294 | Na<sub>2</sub>C<sub>3</sub>H<sub>2</sub>O<sub>3</sub><sup>+</sup> | 174.8890 | Unknown |
| 131.9949 | Na<sub>2</sub>C<sub>3</sub>H<sub>4</sub>NO<sub>2</sub><sup>+</sup> | 175.0912 | Na<sub>2</sub>C<sub>5</sub>H<sub>5</sub>O<sub>4</sub><sup>+</sup> |
| 132.9766 | Na<sub>2</sub>C<sub>3</sub>H<sub>3</sub>O<sub>3</sub><sup>+</sup> | 176.0339 | NaC<sub>6</sub>HO<sub>5</sub><sup>+</sup> |
| 142.0655 | Na<sub>2</sub>C<sub>5</sub>H<sub>4</sub>O<sub>2</sub><sup>+</sup> | 178.9297 | K<sub>2</sub>C<sub>3</sub>HO<sub>4</sub><sup>+</sup> |
| 143.9852 | Na<sub>3</sub>C<sub>2</sub>H<sub>5</sub>NO<sub>2</sub><sup>+</sup> | 179.0615 | Na<sub>3</sub>C<sub>5</sub>H<sub>2</sub>O<sub>3</sub><sup>+</sup> |
| 144.0702 | C<sub>6</sub>H<sub>10</sub>NO<sub>3</sub><sup>+</sup> | 183.0912 | Na<sub>2</sub>C<sub>7</sub>H<sub>5</sub>O<sub>3</sub><sup>+</sup> |
| 146.9881 | Na<sub>2</sub>C<sub>4</sub>H<sub>5</sub>O<sub>3</sub><sup>+</sup> | 186.0848 | NaC<sub>5</sub>H<sub>7</sub>O<sub>6</sub><sup>+</sup> |
| 147.0197 | Na<sub>2</sub>C<sub>4</sub>H<sub>5</sub>O<sub>3</sub><sup>+</sup> | 187.9005 | NaC<sub>5</sub>H<sub>9</sub>O<sub>6</sub><sup>+</sup> |
| 147.9119 | Na<sub>2</sub>C<sub>4</sub>H<sub>5</sub>O<sub>3</sub><sup>+</sup> | 188.8965 | Unknown |
| 152.0382 | Na<sub>3</sub>C<sub>3</sub>HNO<sub>2</sub><sup>+</sup> | 189.8918 | Na<sub>2</sub>C<sub>4</sub>H<sub>2</sub>NO<sub>5</sub><sup>+</sup> |
| 152.9485 | Unknown | 193.0816 | K<sub>2</sub>C<sub>6</sub>H<sub>11</sub>O<sub>2</sub><sup>+</sup> |
| 153.0371 | Na<sub>3</sub>C<sub>4</sub>H<sub>4</sub>O<sub>2</sub><sup>+</sup> | 195.9752 | Na<sub>3</sub>C<sub>5</sub>H<sub>3</sub>O<sub>4</sub><sup>+</sup> |
| 153.0590 | Na<sub>3</sub>C<sub>4</sub>H<sub>4</sub>O<sub>2</sub><sup>+</sup> | 196.1088 | C<sub>7</sub>H<sub>16</sub>O<sub>6</sub><sup>+</sup> |
| 154.0374 | Na<sub>2</sub>C<sub>7</sub>H<sub>8</sub>O<sup>+</sup> | 198.0921 | NaC<sub>6</sub>H<sub>7</sub>O<sub>6</sub><sup>+</sup> |
| 154.0529 | Unknown | 202.8922 | Unknown |
| 156.0775 | NaC<sub>4</sub>H<sub>5</sub>O<sub>5</sub><sup>+</sup> | 204.8912 | Unknown |
| 161.0070 | Na<sub>2</sub>C<sub>5</sub>H<sub>7</sub>O<sub>3</sub><sup>+</sup> | 206.9159 | Unknown |
| 161.8840 | K<sub>2</sub>C<sub>5</sub>H<sub>8</sub>O<sup>+</sup> | 211.1195 | Na<sub>2</sub>C<sub>8</sub>H<sub>5</sub>O<sub>4</sub><sup>+</sup> |
| 162.9370 | Unknown | 213.9199 | Unknown |
| 162.9599 | Na<sub>3</sub>C<sub>4</sub>NO<sub>2</sub><sup>+</sup> | 242.8466 | Unknown |

**Supplementary Table 3.** Negative common GAG ions identified based on their presence in the reference spectra of hyaluronan and at least one sulphated GAG and WT hiPSCs.

| m/z | Assignment | m/z | Assignment |
| --- | --- | --- | --- |
| 81.9917 | Unknown | 134.9219 | Unknown |
| 95.9612 | KC<sub>2</sub>HO<sub>2</sub><sup>-</sup> | 135.9162 | K<sub>2</sub>CNO<sub>2</sub><sup>-</sup> |
| 96.0084 | C<sub>4</sub>H<sub>2</sub>NO<sub>2</sub><sup>-</sup> | 136.0049 | C<sub>6</sub>H<sub>2</sub>NO<sub>3</sub><sup>-</sup> |
| 96.9741 | KC<sub>3</sub>H<sub>6</sub>O<sup>-</sup> | 138.0130 | NaC<sub>4</sub>H<sub>5</sub>NO<sub>3</sub><sup>-</sup> |
| 97.9598 | Unknown | 138.9811 | KC<sub>4</sub>H<sub>4</sub>O<sub>3</sub><sup>-</sup> |
| 98.0206 | C<sub>4</sub>H<sub>4</sub>NO<sub>2</sub><sup>-</sup> | 140.0315 | NaC<sub>4</sub>H<sub>7</sub>NO<sub>3</sub><sup>-</sup> |
| 100.0399 | C<sub>4</sub>H<sub>6</sub>NO<sub>2</sub><sup>-</sup> | 142.9668 | Unknown |
| 104.0343 | Unknown | 148.9198 | KC<sub>4</sub>NO<sub>3</sub><sup>-</sup> |
| 105.9229 | Unknown | 149.9327 | Unknown |
| 105.9828 | Unknown | 149.9944 | K<sub>2</sub>C<sub>4</sub>H<sub>8</sub>O<sup>-</sup> |
| 109.0420 | NaC<sub>4</sub>H<sub>6</sub>O<sub>2</sub><sup>-</sup> | 151.0073 | Unknown |
| 109.9731 | NaC<sub>2</sub>HNO<sub>3</sub><sup>-</sup> | 151.9361 | Unknown |
| 110.0260 | C<sub>5</sub>H<sub>4</sub>NO<sub>2</sub><sup>-</sup> | 152.0260 | NaC<sub>5</sub>H<sub>7</sub>NO<sub>3</sub><sup>-</sup> |
| 111.0117 | C<sub>5</sub>H<sub>3</sub>O<sub>3</sub><sup>-</sup> | 154.0617 | C<sub>7</sub>H<sub>8</sub>NO<sub>3</sub><sup>-</sup> |
| 112.0364 | C<sub>5</sub>H<sub>6</sub>NO<sub>2</sub><sup>-</sup> | 161.9368 | Unknown |
| 112.9633 | Unknown | 163.0218 | Unknown |
| 113.0190 | C<sub>5</sub>H<sub>5</sub>O<sub>3</sub><sup>-</sup> | 163.9152 | NaC<sub>5</sub>HO<sub>5</sub><sup>-</sup> |
| 113.9724 | C<sub>4</sub>H<sub>2</sub>O<sub>4</sub><sup>-</sup> | 163.9391 | NaC<sub>5</sub>HO<sub>5</sub><sup>-</sup> |
| 114.0156 | C<sub>4</sub>H<sub>4</sub>NO<sub>3</sub><sup>-</sup> | 164.9132 | Unknown |
| 117.9182 | Unknown | 166.9473 | Unknown |
| 119.0350 | Na<sub>2</sub>C<sub>3</sub>H<sub>7</sub>NO<sup>-</sup> | 167.9424 | Unknown |
| 119.9302 | Unknown | 168.0600 | C<sub>7</sub>H<sub>6</sub>NO<sub>4</sub><sup>-</sup> |
| 119.9810 | Na<sub>2</sub>C<sub>2</sub>H<sub>2</sub>O<sub>3</sub><sup>-</sup> | 168.9782 | Unknown |
| 121.9102 | K<sub>2</sub>C<sub>2</sub>H<sub>4</sub>O<sup>-</sup> | 171.9884 | Unknown |
| 121.9337 | K<sub>2</sub>C<sub>2</sub>H<sub>4</sub>O<sup>-</sup> | 172.9867 | Na<sub>2</sub>C<sub>5</sub>H<sub>3</sub>O<sub>4</sub><sup>-</sup> |
| 124.0100 | Unknown | 174.0510 | C<sub>7</sub>H<sub>10</sub>O<sub>5</sub><sup>-</sup> |
| 125.9553 | Unknown | 175.8844 | Unknown |
| 126.9628 | KC<sub>2</sub>H<sub>2</sub>NO<sub>3</sub><sup>-</sup> | 176.0336 | C<sub>6</sub>H<sub>8</sub>O<sub>6</sub><sup>-</sup> |
| 131.9678 | Unknown | 176.0568 | C<sub>6</sub>H<sub>8</sub>O<sub>6</sub><sup>-</sup> |
| 132.0302 | Na<sub>2</sub>C<sub>5</sub>H<sub>10</sub>O<sup>-</sup> | 178.0632 | C<sub>6</sub>H<sub>10</sub>O<sub>6</sub><sup>-</sup> |
| 133.0505 | Na<sub>2</sub>C<sub>4</sub>H<sub>9</sub>NO<sup>-</sup> | 179.0309 | NaC<sub>7</sub>H<sub>8</sub>O<sub>4</sub><sup>-</sup> |
| 133.9265 | Unknown | 179.9131 | Unknown |
| 133.9822 | NaC<sub>4</sub>HNO<sub>3</sub><sup>-</sup> | 181.9271 | Unknown |
| 134.0488 | Na<sub>2</sub>C<sub>4</sub>H<sub>8</sub>O<sub>2</sub><sup>-</sup> | 185.0131 | Unknown |

**Supplementary Table 4.**
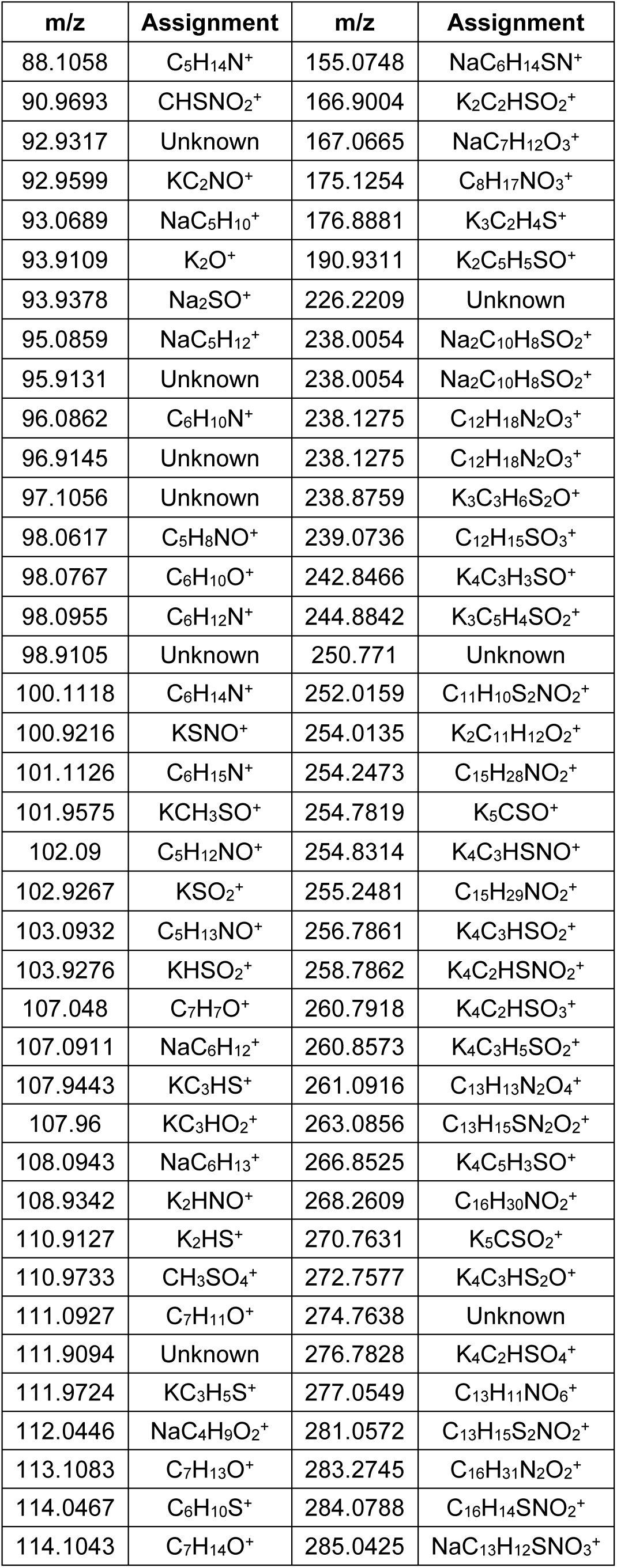
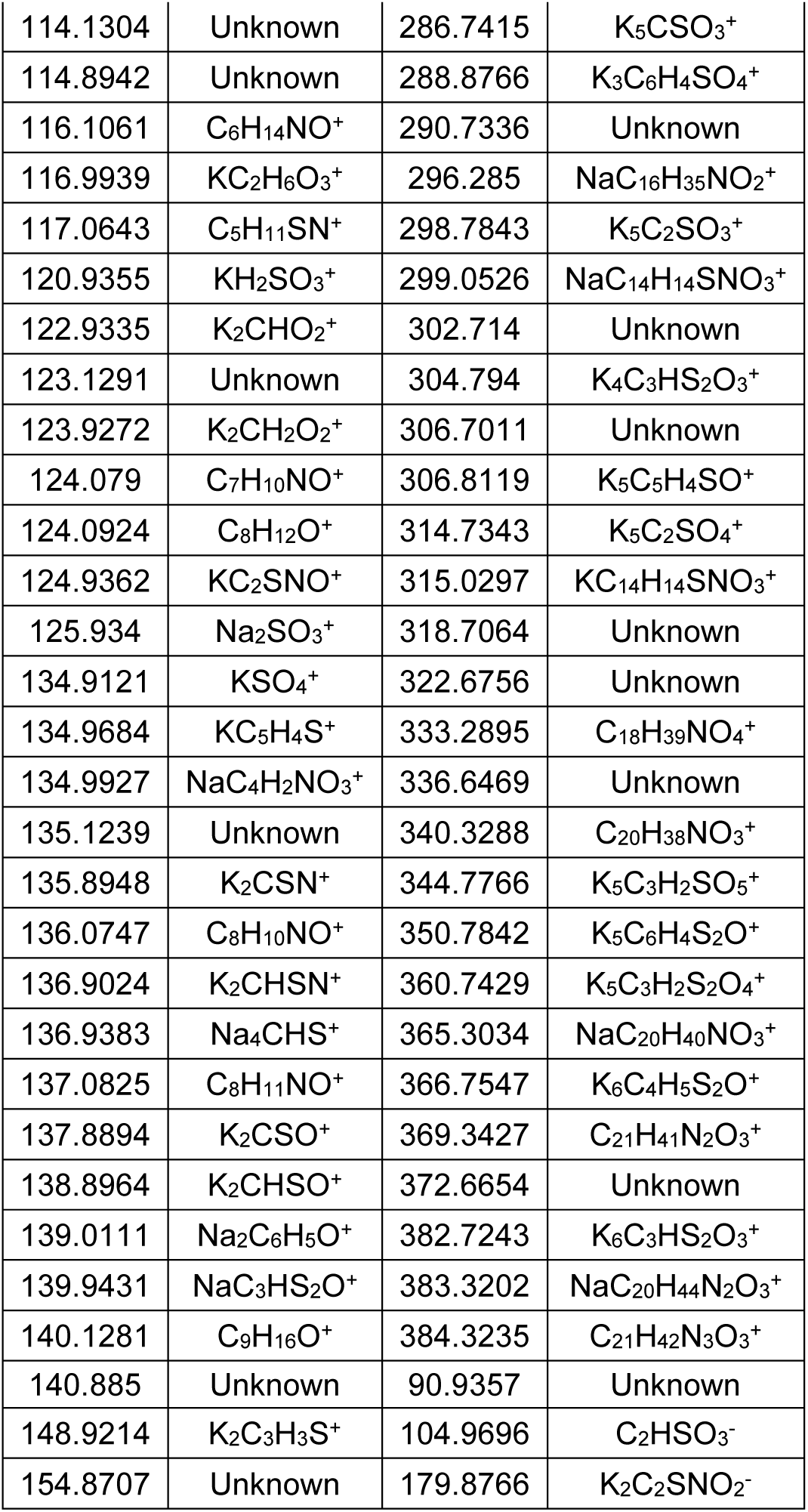
Putative HS ions identified through presence exclusively in HS reference spectra and detection in WT hiPSC.

**Supplementary Table 5.** Putative CS ions identified through presence exclusively in CS reference spectra and detection in WT hiPSCs.

| m/z | Assignment | m/z | Assignment |
| --- | --- | --- | --- |
| 104.0585 | NaC<sub>6</sub>H<sub>9</sub><sup>+</sup> | 239.124 | C<sub>12</sub>H<sub>17</sub>NO<sub>4</sub><sup>-</sup> |
| 104.0998 | C<sub>5</sub>H<sub>14</sub>NO<sup>+</sup> | 239.2256 | C<sub>14</sub>H<sub>25</sub>NO<sub>2</sub><sup>-</sup> |
| 127.0909 | C<sub>7</sub>H<sub>13</sub>NO<sup>+</sup> | 240.026 | C<sub>11</sub>H<sub>12</sub>S<sub>2</sub>O<sub>2</sub><sup>-</sup> |
| 186.8886 | K<sub>2</sub>CHSO<sub>4</sub><sup>+</sup> | 240.1194 | C<sub>12</sub>H<sub>18</sub>NO<sub>4</sub><sup>-</sup> |
| 327.3339 | C<sub>19</sub>H<sub>37</sub>NO<sub>3</sub><sup>+</sup> | 241.0569 | KC<sub>12</sub>H<sub>12</sub>NO<sub>2</sub><sup>-</sup> |
| 194.872 | K<sub>2</sub>C<sub>3</sub>HS<sub>2</sub>O<sup>-</sup> | 242.7733 | Unknown |
| 194.8988 | K<sub>2</sub>C<sub>3</sub>H<sub>3</sub>S<sub>2</sub>N<sup>-</sup> | 242.9026 | KC<sub>6</sub>H<sub>4</sub>S<sub>3</sub>O<sub>2</sub><sup>-</sup> |
| 195.0337 | C<sub>9</sub>H<sub>9</sub>SNO<sub>2</sub><sup>-</sup> | 244.8705 | K<sub>3</sub>C<sub>4</sub>H<sub>2</sub>SNO<sub>2</sub><sup>-</sup> |
| 195.9266 | Na<sub>2</sub>C<sub>3</sub>H<sub>2</sub>S<sub>2</sub>O<sub>3</sub><sup>-</sup> | 244.9191 | KC<sub>6</sub>H<sub>6</sub>S<sub>3</sub>O<sub>2</sub><sup>-</sup> |
| 195.9499 | Na<sub>3</sub>C<sub>5</sub>H<sub>3</sub>SO<sub>2</sub><sup>-</sup> | 246.0629 | C<sub>12</sub>H<sub>10</sub>N<sub>2</sub>O<sub>4</sub><sup>-</sup> |
| 196.0518 | NaC<sub>11</sub>H<sub>9</sub>O<sub>2</sub><sup>-</sup> | 254.0027 | C<sub>11</sub>H<sub>10</sub>S<sub>2</sub>O<sub>3</sub><sup>-</sup> |
| 196.8752 | K<sub>2</sub>C<sub>3</sub>H<sub>3</sub>S<sub>2</sub>O<sup>-</sup> | 255.239 | C<sub>15</sub>H<sub>29</sub>NO<sub>2</sub><sup>-</sup> |
| 196.9223 | C<sub>3</sub>HS<sub>2</sub>O<sub>6</sub><sup>-</sup> | 256.2421 | C<sub>15</sub>H<sub>30</sub>NO<sub>2</sub><sup>-</sup> |
| 197.9337 | KC<sub>5</sub>H<sub>3</sub>SO<sub>4</sub><sup>-</sup> | 256.2724 | Unknown |
| 198.9155 | KC<sub>4</sub>H<sub>2</sub>S<sub>2</sub>NO<sub>2</sub><sup>-</sup> | 257.2463 | C<sub>15</sub>H<sub>31</sub>NO<sub>2</sub><sup>-</sup> |
| 199.9049 | K<sub>3</sub>C<sub>4</sub>H<sub>3</sub>O<sub>2</sub><sup>-</sup> | 258.8388 | K<sub>4</sub>C<sub>3</sub>H<sub>3</sub>SO<sub>2</sub><sup>-</sup> |
| 199.9412 | KC<sub>5</sub>H<sub>5</sub>S<sub>2</sub>O<sub>2</sub><sup>-</sup> | 259.0991 | KC<sub>13</sub>H<sub>18</sub>NO<sub>2</sub><sup>-</sup> |
| 201.9918 | C<sub>7</sub>H<sub>6</sub>SO<sub>5</sub><sup>-</sup> | 260.9073 | K<sub>2</sub>C<sub>7</sub>H<sub>5</sub>S<sub>2</sub>NO<sup>-</sup> |
| 202.8624 | K<sub>3</sub>C<sub>2</sub>SNO<sup>-</sup> | 261.8531 | K<sub>3</sub>C<sub>4</sub>H<sub>3</sub>S<sub>2</sub>NO<sup>-</sup> |
| 203.0843 | KC<sub>11</sub>H<sub>16</sub>O<sup>-</sup> | 262.8618 | K<sub>3</sub>C<sub>4</sub>H<sub>2</sub>SO<sub>4</sub><sup>-</sup> |
| 203.8837 | K<sub>3</sub>C<sub>3</sub>H<sub>3</sub>SO<sup>-</sup> | 262.8931 | K<sub>3</sub>C<sub>5</sub>H<sub>6</sub>SO<sub>3</sub><sup>-</sup> |
| 204.9069 | K<sub>3</sub>C<sub>3</sub>H<sub>4</sub>O<sub>3</sub><sup>-</sup> | 264.8596 | K<sub>3</sub>C<sub>4</sub>H<sub>4</sub>S<sub>2</sub>O<sub>2</sub><sup>-</sup> |
| 206.9016 | K<sub>2</sub>C<sub>4</sub>H<sub>3</sub>S<sub>2</sub>N<sup>-</sup> | 265.2452 | C<sub>16</sub>H<sub>27</sub>NO<sub>2</sub><sup>-</sup> |
| 207.8968 | K<sub>2</sub>C<sub>4</sub>H<sub>2</sub>SO<sub>3</sub><sup>-</sup> | 266.0262 | C<sub>12</sub>H<sub>10</sub>SO<sub>5</sub><sup>-</sup> |
| 208.0772 | C<sub>11</sub>H<sub>14</sub>SNO<sup>-</sup> | 266.7938 | K<sub>5</sub>C<sub>2</sub>SO<sup>-</sup> |
| 208.1021 | C<sub>11</sub>H<sub>14</sub>NO<sub>3</sub><sup>-</sup> | 266.848 | K<sub>4</sub>C<sub>5</sub>H<sub>3</sub>SO<sup>-</sup> |
| 208.8737 | K<sub>2</sub>C<sub>3</sub>HS<sub>2</sub>NO<sup>-</sup> | 268.8266 | K<sub>3</sub>C<sub>2</sub>SO<sub>6</sub><sup>-</sup> |
| 209.0571 | Na<sub>2</sub>C<sub>10</sub>H<sub>11</sub>O<sub>2</sub><sup>-</sup> | 270.8385 | K<sub>4</sub>C<sub>4</sub>H<sub>3</sub>SO<sub>2</sub><sup>-</sup> |
| 210.0571 | NaC<sub>11</sub>H<sub>9</sub>NO<sub>2</sub><sup>-</sup> | 271.2235 | C<sub>15</sub>H<sub>29</sub>NO<sub>3</sub><sup>-</sup> |
| 210.8421 | K<sub>2</sub>C<sub>3</sub>HS<sub>2</sub>O<sub>2</sub><sup>-</sup> | 272.8405 | K<sub>4</sub>C<sub>3</sub>H<sub>3</sub>S<sub>2</sub>N<sup>-</sup> |
| 210.8717 | K<sub>2</sub>C<sub>3</sub>HS<sub>2</sub>O<sub>2</sub><sup>-</sup> | 276.832 | K<sub>3</sub>C<sub>4</sub>H<sub>2</sub>S<sub>2</sub>NO<sub>2</sub><sup>-</sup> |
| 211.9134 | K<sub>2</sub>C<sub>4</sub>H<sub>6</sub>S<sub>2</sub>O<sup>-</sup> | 276.8951 | K<sub>3</sub>C<sub>6</sub>H<sub>8</sub>S<sub>2</sub>O<sup>-</sup> |
| 211.9465 | KC<sub>6</sub>H<sub>5</sub>SO<sub>4</sub><sup>-</sup> | 278.8692 | K<sub>4</sub>C<sub>6</sub>H<sub>5</sub>SN<sup>-</sup> |
| 212.8304 | K<sub>4</sub>C<sub>2</sub>HS<sup>-</sup> | 279.2562 | NaC<sub>15</sub>H<sub>30</sub>NO<sub>2</sub><sup>-</sup> |
| 212.866 | K<sub>3</sub>C<sub>4</sub>H<sub>2</sub>SN<sup>-</sup> | 279.9228 | K<sub>3</sub>C<sub>9</sub>H<sub>7</sub>O<sub>3</sub><sup>-</sup> |
| 213.8696 | $\text{K}_2\text{C}_2\text{SO}_5^-$ | 280.2586 | $\text{NaC}_{15}\text{H}_{31}\text{NO}_2^-$ |
| 214.8857 | $\text{K}_2\text{C}_2\text{HSO}_5^-$ | 281.2704 | $\text{C}_{16}\text{H}_{29}\text{N}_2\text{O}_2^-$ |
| 216.9437 | $\text{C}_7\text{H}_5\text{S}_3\text{O}_2^-$ | 282.2948 | Unknown |
| 219.8686 | $\text{K}_3\text{C}_2\text{HSNO}_2^-$ | 282.7534 | $\text{K}_5\text{C}_2\text{SO}_2^-$ |
| 220.1261 | $\text{C}_{12}\text{H}_{16}\text{N}_2\text{O}_2^-$ | 282.8652 | $\text{K}_4\text{C}_5\text{H}_3\text{O}_4^-$ |
| 220.9015 | $\text{K}_2\text{C}_5\text{H}_3\text{SO}_3^-$ | 283.2782 | $\text{C}_{16}\text{H}_{31}\text{N}_2\text{O}_2^-$ |
| 222.0941 | $\text{C}_{12}\text{H}_{14}\text{O}_4^-$ | 284.2778 | $\text{C}_{16}\text{H}_{32}\text{N}_2\text{O}_2^-$ |
| 223.0605 | $\text{C}_{11}\text{H}_{11}\text{O}_5^-$ | 296.8295 | $\text{K}_4\text{C}_5\text{H}_3\text{S}_2\text{N}^-$ |
| 223.1834 | $\text{NaC}_{11}\text{H}_{24}\text{N}_2\text{O}^-$ | 299.0694 | $\text{Na}_2\text{C}_{15}\text{H}_{13}\text{N}_2\text{O}_2^-$ |
| 224.0915 | $\text{C}_{11}\text{H}_{14}\text{NO}_4^-$ | 299.1306 | $\text{KC}_{16}\text{H}_{22}\text{NO}_2^-$ |
| 224.804 | $\text{K}_4\text{C}_3\text{HS}^-$ | 300.7222 | Unknown |
| 224.9244 | $\text{K}_2\text{C}_5\text{H}_7\text{S}_2\text{O}^-$ | 300.8381 | $\text{K}_4\text{C}_5\text{H}_5\text{S}_2\text{O}^-$ |
| 225.892 | $\text{K}_2\text{C}_4\text{H}_4\text{S}_2\text{O}_2^-$ | 300.8822 | $\text{K}_3\text{C}_8\text{H}_8\text{S}_2\text{O}^-$ |
| 225.9968 | $\text{K}_2\text{C}_9\text{H}_{10}\text{NO}^-$ | 302.7209 | Unknown |
| 227.2144 | $\text{C}_{13}\text{H}_{25}\text{NO}_2^-$ | 302.8543 | $\text{K}_4\text{C}_5\text{H}_7\text{S}_2\text{O}^-$ |
| 227.9998 | $\text{KC}_{11}\text{H}_9\text{SO}^-$ | 305.2651 | $\text{C}_{15}\text{H}_{33}\text{N}_2\text{O}_4^-$ |
| 228.2075 | $\text{C}_{13}\text{H}_{26}\text{NO}_2^-$ | 308.2857 | $\text{NaC}_{16}\text{H}_{33}\text{N}_2\text{O}_2^-$ |
| 233.0756 | $\text{C}_{12}\text{H}_{11}\text{NO}_4^-$ | 309.321 | Unknown |
| 233.116 | $\text{NaC}_{12}\text{H}_{18}\text{O}_3^-$ | 311.1556 | $\text{C}_{16}\text{H}_{25}\text{SNO}_3^-$ |
| 234.8624 | $\text{K}_3\text{C}_3\text{H}_2\text{SO}_3^-$ | 312.179 | $\text{C}_{16}\text{H}_{26}\text{NO}_5^-$ |
| 234.9034 | $\text{K}_2\text{C}_6\text{H}_5\text{S}_2\text{O}^-$ | 315.0632 | $\text{KC}_{15}\text{H}_{18}\text{SNO}_2^-$ |
| 236.0867 | $\text{NaC}_{11}\text{H}_{17}\text{SO}_2^-$ | 322.8817 | $\text{K}_2\text{C}_9\text{H}_9\text{S}_3\text{O}_2^-$ |
| 236.8619 | $\text{K}_3\text{C}_3\text{H}_4\text{S}_2\text{O}^-$ | 328.728 | $\text{K}_5\text{C}_3\text{H}_2\text{S}_2\text{O}_2^-$ |
| 237.0944 | $\text{C}_{12}\text{H}_{15}\text{NO}_4^-$ | 328.8152 | $\text{K}_4\text{C}_5\text{H}_3\text{S}_2\text{NO}_2^-$ |
| 238.1324 | $\text{C}_{12}\text{H}_{18}\text{N}_2\text{O}_3^-$ | | |

**Supplementary Table 6:** Putative DS ions identified through presence exclusively in DS reference spectra and detection in WT hiPSCs.

| m/z | Assignment |
| --- | --- |
| 146.0181 | $C_5H_6O_5^+$ |
| 146.1041 | $C_7H_{14}O_3^+$ |
| 164.0255 | $NaC_7H_9SO^+$ |
| 244.8404 | $K_4C_2H_3SNO^+$ |
| 82.0277 | $NaC_2H_5NO^-$ |
| 87.96438 | $C_2SO_2^-$ |
| 88.01885 | $NaC_4H_3N^-$ |
| 115.9556 | $Na_2C_2SN^-$ |
| 116.0533 | $C_5H_{10}SN^-$ |
| 140.0456 | $NaC_5H_9O_3^-$ |
| 140.0618 | $NaC_5H_{11}NO_2^-$ |

**Supplementary Table 7:** Putative KS ions identified through presence exclusively in KS reference spectra and detection in WT hiPSCs.

| m/z | Assignment |
| --- | --- |
| 106.0646 | $NaC_5H_9N^+$ |
| 106.0752 | $NaC_6H_{11}^+$ |
| 82.0712 | $C_5H_8N^-$ |
| 84.97157 | $NaCH_2SO^-$ |

**Supplementary Table 8:** Putative HA ions identified through presence exclusively in HA reference spectra and detection in WT hiPSCs.

| m/z | Assignment |
| --- | --- |
| 187.9915 | $Na_3C_3H_5NO_4^-$ |
| 189.0547 | $C_{11}H_9O_3^-$ |
| 190.0433 | $KC_9H_{11}O_2^-$ |
| 191.0752 | $KC_9H_{14}NO^-$ |
| 191.8909 | Unknown |
| 192.0715 | $C_{10}H_{10}NO_3^-$ |
| 192.8679 | Unknown |
| 193.0393 | $C_9H_7NO_4^-$ |
| 193.0592 | $Na_2C_{10}H_{11}O^-$ |
| 193.0808 | $NaC_9H_{14}O_3^-$ |
| 193.9822 | $Na_3C_5H_3NO_3^-$ |

## Bibliography

1. Merry, C. L. R., Lindahl, U., Couchman, J. & Esko, J. D. Proteoglycans and Sulfated Glycosaminoglycans. (2022).

2. Garcí;a-García, M. J. & Anderson, K. V. Essential Role of Glycosaminoglycans in Fgf Signaling during Mouse Gastrulation. Cell 114, 727–737 (2003).

3. Betteridge, K. B. et al. Sialic acids regulate microvessel permeability, revealed by novel *in vivo* studies of endothelial glycocalyx structure and function. J. Physiol. 595, 5015–5035 (2017).

4. Wei, J., Hu, M., Huang, K., Lin, S. & Du, H. Roles of Proteoglycans and Glycosaminoglycans in Cancer Development and Progression. Int. J. Mol. Sci. 21, 5983 (2020).

5. Gamez, M. et al. Heparanase inhibition as a systemic approach to protect the endothelial glycocalyx and prevent microvascular complications in diabetes. Cardiovasc. Diabetol. 23, 50 (2024).

6. Sullivan, R. C., Rockstrom, M. D., Schmidt, E. P. & Hippensteel, J. A. Endothelial glycocalyx degradation during sepsis: Causes and consequences. Matrix Biol. Plus 12, 100094 (2021).

7. Milne, L. K. et al. Validation of the use of ToF-SIMS for analysis of glycosaminoglycans. Carbohydr. Polym. 371, 124415 (2026).

8. Qiu, Y. et al. Endothelial glycocalyx is damaged in diabetic cardiomyopathy: angiopoietin 1 restores glycocalyx and improves diastolic function in mice. Diabetologia 65, 879–894 (2022).

9. Davies-Strickleton, H., et al. Collection of protocols for label-free GAG disaccharide analysis by HILIC-MS/MS of diverse biological tissues v1. Preprint at 10.17504/protocols.io.x54v956n1l3e/v1 (2025).

10. Basu, A. et al. Quantitative HILIC-Q-TOF-MS Analysis of Glycosaminoglycans and Non-reducing End Carbohydrate Biomarkers via Glycan Reductive Isotopic Labeling. Anal. Chem. 97, 17490–17500 (2025).

11. Shao, C., Shi, X., Phillips, J. J. & Zaia, J. Mass Spectral Profiling of Glycosaminoglycans from Histological Tissue Surfaces. Anal. Chem. 85, 10984–10991 (2013).

12. Przybylski, C., Gonnet, F., Buchmann, W. & Daniel, R. Critical parameters for the analysis of anionic oligosaccharides by desorption electrospray ionization mass spectrometry. Journal of Mass Spectrometry 47, 1047–1058 (2012).

13. Palomino, T. V. & Muddiman, D. C. Glycosaminoglycan Mass Spectrometry Imaging by Infrared Matrix-Assisted Laser Desorption Electrospray Ionization. J. Am. Soc. Mass Spectrom. 36, 658–663 (2025).

14. Passarelli, M. K. et al. The 3D OrbiSIMS—label-free metabolic imaging with subcellular lateral resolution and high mass-resolving power. Nat. Methods 14, 1175–1183 (2017).

15. Passarelli, M. K. & Winograd, N. Lipid imaging with time-of-flight secondary ion mass spectrometry (ToF-SIMS). Biochimica et Biophysica Acta (BBA) - Molecular and Cell Biology of Lipids 1811, 976–990 (2011).

16. Kotowska, A. M. et al. Protein identification by 3D OrbiSIMS to facilitate in situ imaging and depth profiling. Nat. Commun. 11, 5832 (2020).

17. Ward, S. et al. Integrating cryo-OrbiSIMS with computational modelling and metadynamics simulations enhances RNA structure prediction at atomic resolution. Nat. Commun. 15, 4367 (2024).

18. Kubicek, M. et al. A novel ToF-SIMS operation mode for sub 100nm lateral resolution: Application and performance. Appl. Surf. Sci. 289, 407–416 (2014).

19. Suvannapruk, W. et al. Single-Cell Metabolic Profiling of Macrophages Using 3D OrbiSIMS: Correlations with Phenotype. Anal. Chem. 94, 9389–9398 (2022).

20. Hook, A. L., Hogwood, J., Gray, E., Mulloy, B. & Merry, C. L. R. High sensitivity analysis of nanogram quantities of glycosaminoglycans using ToF-SIMS. Commun. Chem. 4, 67 (2021).

21. Linke, F. et al. Identifying new biomarkers of aggressive Group 3 and SHH medulloblastoma using 3D hydrogel models, single cell RNA sequencing and 3D OrbiSIMS imaging. Acta Neuropathol. Commun. 11, 6 (2023).

22. Zimmermann, R. et al. Discriminant Principal Component Analysis of ToF-SIMS Spectra for Deciphering Compositional Differences of MSC-Secreted Extracellular Matrices. Small Methods 7, (2023).

23. Chen, Y.-H. et al. The GAGOme: a cell-based library of displayed glycosaminoglycans. Nat. Methods 15, 881–888 (2018).

24. Karlsson, R. et al. Dissecting structure-function of 3-O-sulfated heparin and engineered heparan sulfates. Sci. Adv. 7, (2021).

25. Stickens, D., Zak, B. M., Rougier, N., Esko, J. D. & Werb, Z. Mice deficient in Ext2 lack heparan sulfate and develop exostoses. Development 132, 5055–5068 (2005).

26. Syangtan, D. et al. Heparan sulfate regulates the fate decisions of human pluripotent stem cells. Stem Cell Reports 20, 102384 (2025).

27. Pareek, V., Tian, H., Winograd, N. & Benkovic, S. J. Metabolomics and mass spectrometry imaging reveal channeled de novo purine synthesis in cells. Science (1979). 368, 283–290 (2020).

28. Wilson, R. L. et al. Fluorinated Colloidal Gold Immunolabels for Imaging Select Proteins in Parallel with Lipids Using High-Resolution Secondary Ion Mass Spectrometry. Bioconjug. Chem. 23, 450–460 (2012).

29. Van Ham, R., Van Vaeck, L., Adams, F. C. & Adriaens, A. Systematization of the Mass Spectra for Speciation of Inorganic Salts with Static Secondary Ion Mass Spectrometry. Anal. Chem. 76, 2609–2617 (2004).

30. Ramnath, R. D. et al. Blocking matrix metalloproteinase-mediated syndecan-4 shedding restores the endothelial glycocalyx and glomerular filtration barrier function in early diabetic kidney disease. Kidney Int. 97, 951–965 (2020).

31. Oltean, S. et al. Vascular Endothelial Growth Factor-A165b Is Protective and Restores Endothelial Glycocalyx in Diabetic Nephropathy. Journal of the American Society of Nephrology 26, 1889–1904 (2015).

32. van den Hoven, M. J., et al. Increased expression of heparanase in overt diabetic nephropathy. Kidney Int. 70, 2100–2108 (2006).

33. Shi, X. et al. A ToF-SIMS methodology for analyzing inter-tissue lipid distribution variations and intra-tissue multilevel mass spectrometry imaging within a single rat. Microchemical Journal 200, 110235 (2024).

34. Gamble, L. J. et al. ToF-SIMS of tissues: “Lessons learned” from mice and women. Biointerphases 10, (2015).

35. Wu, L. et al. Imaging and differentiation of mouse embryo tissues by ToF-SIMS. Int. J. Mass Spectrom. 260, 137–145 (2007).

36. Deakin, J. A. & Lyon, M. A simplified and sensitive fluorescent method for disaccharide analysis of both heparan sulfate and chondroitin/dermatan sulfates from biological samples. Glycobiology 18, 483–491 (2008).

37. Song, W. et al. The lipoprotein lipase that is shuttled into capillaries by GPIHBP1 enters the glycocalyx where it mediates lipoprotein processing. Proceedings of the National Academy of Sciences 120, (2023).

38. Winograd, N. & Bloom, A. Sample Preparation for 3D SIMS Chemical Imaging of Cells. in 9–19 (2015). doi:10.1007/978-1-4939-1357-2_2.

39. Mosqueira, D. et al. CRISPR/Cas9 editing in human pluripotent stem cell-cardiomyocytes highlights arrhythmias, hypocontractility, and energy depletion as potential therapeutic targets for hypertrophic cardiomyopathy. Eur. Heart J. 39, 3879–3892 (2018).

40. Kondrashov, A. et al. Simplified Footprint-Free Cas9/CRISPR Editing of Cardiac-Associated Genes in Human Pluripotent Stem Cells. Stem Cells Dev. 27, 391–404 (2018).

41. Watson, H. A. et al. Heparan Sulfate Inhibits Hematopoietic Stem and Progenitor Cell Migration and Engraftment in Mucopolysaccharidosis I. Journal of Biological Chemistry 289, 36194–36203 (2014).

42. Holley, R. J. et al. Use of Flow Cytometry for Characterization and Fractionation of Cell Populations Based on Their Expression of Heparan Sulfate Epitopes. in 239–251 (2015). doi:10.1007/978-1-4939-1714-3_21.

43. Holley, R. J. et al. Mucopolysaccharidosis Type I, Unique Structure of Accumulated Heparan Sulfate and Increased N-Sulfotransferase Activity in Mice Lacking α-l-iduronidase. Journal of Biological Chemistry 286, 37515–37524 (2011).

44. Dagälv, A. et al. Heparan sulfate structure: methods to study N-sulfation and NDST action. Methods Mol. Biol. 1229, 189–200 (2015).

45. David, G., Bai, X. M., Van der Schueren, B., Cassiman, J. J. & Van den Berghe, H. Developmental changes in heparan sulfate expression: in situ detection with mAbs. J. Cell Biol. 119, 961–75 (1992).

